# Integrating lineage-specific and universal genomic probes illuminates phylogenetic relationships and molecular evolution in Sauvagesieae (Ochnaceae)

**DOI:** 10.64898/2026.04.29.721621

**Authors:** Sandra Reinales, Félix Forest, Alexandre R. Zuntini, Domingos Cardoso, Gustavo A. Ballen, Dairon Cárdenas, José R. Pirani

## Abstract

Obtaining large and well-resolved phylogenetic trees for neotropical clades is challenging, as many species inhabit remote regions, and sampling often relies on herbarium specimens with highly degraded DNA. Target capture provides an effective solution for retrieving molecular data from fragmentary material. However, data processing using tools generally designed for diploid organisms and single-copy loci is also challenging, particularly when events such as genome duplication and hybridisation have shaped the lineage evolution. We used dual-hybridisation to integrate Ochnaceae-specific and universal probes to reconstruct the phylogenetic relationships of Sauvagesieae, a pantropical clade with *ca.* 90 species mainly distributed in the northern Andes, the Brazilian Espinhaço Range, and the Amazon-Guyana region. We tested different filtering strategies involving missing data and paralogs to assess probable sources of tree discordance and topological uncertainty. We found no significant benefit in reducing tree discordance after removing entire genes due to the presence of paralogs or a high amount of missing data. Removing fragmentary sequences instead improved alignments and increased branch support of gene trees. By quantifying the proportion of SNPs, analysing the distribution of the allele frequencies, and gene-tree quartet frequencies, we found evidence of polyploidisation and hybridisation, which could reduce resolution at internal nodes, particularly in mountain clades. Our results underscore the importance of exploring the complexities of target-capture data, not only to improve phylogenetic resolution but also to understand the sources of phylogenetic conflict and the underlying molecular evolutionary processes.

## 1 Introduction

Despite the ever-growing technologically advanced set of methods to analyse phylogenomic data, obtaining comprehensively sampled and well-resolved phylogenetic trees for neotropical plant groups is still challenging, because many species occur in remote regions and sampling often relies largely on herbarium specimens with highly degraded DNA. Target capture is a valuable solution to recover molecular data from herbarium material to reconstruct phylogenetic trees, usually based on hundreds of nuclear loci while allowing the recovery of plastid regions as by-catch (McKain et al., 2018; Pezzini et al., 2023). However, a key insight from empirical studies applying this technique to build well-sampled phylogenetic trees across plant groups is that missing data and orthology assessment from fragmentary and low-quality sequences is a pervasive issue (Frost et al., 2024; Brewer et al., 2019; Bakker, 2017).

While universal probe sets are effective for different flowering plant families and produce highly compatible results across studies (Johnson et al., 2019), lineage-specific probes are generally designed to incorporate low-copy genes with greater variability for the target group, but conserving a great fidelity between probe and target (Schneider et al., 2021; Siniscalchi et al., 2021). The choice between universal and lineage-specific probe sets can be further complicated when previously generated data are available for a subset of samples. Combining universal and lineage-specific probe sets in a dual-hybridisation reaction allows to simultaneously generate sequences from multiple probe sets with relatively little extra cost and optimises laboratory effort (Hendriks et al., 2021). This approach is helpful for obtaining large datasets that can be combined with data from previous studies to obtain more densely sampled phylogenetic trees.

The availability of hundreds to thousands of unlinked loci allows us to look beyond phylogenetic relationships by exploring evolutionary processes and biases that often appear in phylogenomic analyses as tree discordance. Multiple factors can generate discordance between gene trees and the species tree, including biological processes such as incomplete lineage sorting (ILS), gene duplication/loss, and hybridisation, as well as methodological sources of error including data assembly and filtering issues, gene-tree estimation error (GTEE), and random noise from uninformative genes (Morales-Briones et al., 2021b; Molloy and Warnow, 2017; Pardo-De la Hoz et al., 2023). Whole-genome duplication (WGD) events are common throughout the eukaryotic tree of life, and are particularly significant for the evolution of plants, as 30% to 70% of vascular plant species are estimated to have a polyploid origin (Barker et al., 2016; Landis et al., 2018; Jiao et al., 2011). Malpighiales is one of the recalcitrant nodes in the angiosperms phylogeny showing recurrent tree discordance between nuclear and plastid phylogenetic trees, lack of resolution concerning its phylogenetic position, and weak node support throughout (Zuntini et al., 2024; APG IV, 2016; Xi et al., 2012; Cai et al., 2021). The Malpighiales genome has been shaped by evolutionary processes such as genome duplication (Fawcett et al., 2009; Cai et al., 2019), hybridisation (Marcussen et al., 2015), and horizontal gene transfer (Davis, 2004), or even a combination of these factors (Cai et al., 2021). About 22–24 WGDs have been reported broadly distributed across the Malpighiales phylogeny, some of them commonly associated with the highly diverse clades in the order such as ochnoids, clusioids, euphorbioids, phyllanthoids, Violaceae, and Passifloraceae (Cai et al., 2019). Currently, no method is designed to address all these phenomena simultaneously or to separate their effects on DNA sequences produced by short-read assembly. Consequently, using multiple phylogenetic tools and data-partitioning strategies to analyse phylogenomic data is essential for disentangling the potential sources of gene tree discordance and resolving recalcitrant relationships.

Within Malpighiales, tribe Sauvagesieae of Ochnaceae is a predominantly neotropical clade comprising *ca.* 90 species and 17 genera, with *Sauvagesia* L. and *Tyleria* Gleason encompassing around 60% of the extant diversity of the tribe (Schneider et al., 2020). Most species are in the northernmost Andes, the Brazilian Espinhaço Range, and the Amazon-Guyana region (Reinales and Parra-O., 2020; Queiroz-Lima et al., 2023). Recent efforts to increase species sampling and use genomic data from different sources (*i.e.*, nuclear and plastid genomes) have improved our understanding of relationships within the tribe at the generic level (Schneider et al., 2020). Currently, Sauvagesieae is composed of four strongly supported clades, the *Cespedesia* clade + *Fleurydora* A.Chev., *Wallacea* Spruce ex Benth. & Hook.f. + *Poecilandra* Tul., the Southeast Asian clade, and a clade including *Adenarake* Maguire & Wurdack and the paraphyletic *Sauvagesia* L. However, there are conflicting relationships between concatenated and coalescent-based analyses using nuclear genes, and between plastid and nuclear topologies (Reinales and Parra-O., 2020; Schneider et al., 2014, 2020, 2021). Those conflicts are mainly due to the phylogenetic placement of *Blastemanthus* Planch. and *Neckia* Korth, as well as the lack of resolution and low support (local posterior probability, *LPP <* 0.6) of some internal branches within *Sauvagesia* and the SE Asian clade (Schneider et al., 2020). Interestingly, those branches had high bootstrap support in the tree based on concatenated nuclear genes. However, bootstrap support values could be massively inflated in genomic-scale phylogenetic datasets, even assigning high support for the ‘wrong’ topology when assuming that independent genes have the same evolutionary history (*i.e.*, in a concatenated supermatrix approach) in the presence of ILS (Lanfear and Hahn, 2024; Mirarab, 2019; Minh et al., 2020). The impact of gene tree discordance on the phylogenetic relationships within Ochnaceae, as well as its probable causes remains unexplored.

Aiming to address remaining questions about the phylogenetic relationships within Sauvagesieae, we used the dual-hybridisation approach integrating a specific set of probes designed for Ochnaceae (Schneider et al., 2020), and the universal probes Angiosperms353 (Johnson et al., 2019), and apply them to a rich sample of Sauvagesieae (all genera and 90% of the species). This approach also allows the combination of molecular data previously generated using both probe sets separately (Schneider et al., 2020; Shah et al., 2021) with the new data, to produce the most complete phylogenetic tree for the tribe to date. In addition, obtaining sequences for more loci enabled us to assess different sources of topological incongruence such as hybridisation by inferring phylogenetic networks, and polyploidisation via ploidy estimation. By exploring the impact of multiple filtering strategies on phylogenetic reconstruction, along with the possible effects of reticulate evolution and gene duplication events, we aim to elucidate the evolutionary relationships within Sauvagesieae, thereby illustrating another versatile framework for the analysis of target-capture datasets in tropical plants.

## 2 Materials and methods

### 2.1 Taxon sampling

New DNA sequences were produced for 96 samples and 73 species of Sauvagesieae based on specimens collected in the field or sampled from herbaria (Table S1). Additional raw data produced by Schneider et al. (2020); Shah et al. (2021, 2022); Baker et al. (2022), corresponding to 45 species within Sauvagesieae and 11 species used as outgroup taxa were downloaded from the EMBL European Bioinformatics Institute (Table S1). After combining newly sequenced and already-published data, we analysed 170 samples representing all genera and 80 species, *i.e.*, 90% of the species richness of Sauvagesieae.

### 2.2 DNA extraction and library preparation

Genomic DNA was extracted from up to 20 mg of leaf tissue (silica-dried or herbarium-preserved) using a modified CTAB protocol (Doyle and Doyle, 1987; Tel-Zur et al., 1999, see supplementary materials for details; Fig. S1). Samples with more than 300 ng of DNA were purified with Agencourt AMPure XP Beads (Beckman Coulter, Indianapolis, IN, USA) following the manufacturer’s protocol. All DNA extracts were run on a 1% agarose gel to assess their average fragment size. Samples with low concentration (not visible on a 1% agarose gel) were assessed on an Agilent Technologies 4200 TapeStation System (Santa Clara, CA, USA). DNA extracts with an average fragment size *>* 350 bp were sonicated using a Covaris M220 Focused-ultrasonicator (Covaris, Woburn, MA, USA) following the manufacturer’s protocol.

Dual-indexed libraries for Illumina sequencing were prepared using the NEBNEXT Ultra II DNA Library Prep Kit and the NEBNEXT Multiplex Oligos for Illumina (Dual Index Primers 5 and 7; New England BioLabs, Ipswich, MA, USA) following the manufacturer’s protocol, but using half the recommended volumes and applying size selection depending on the DNA input. DNA fragments were amplified using 5-8(−12) cycles of PCR. Libraries were quantified using a Quantus fluorometer, and fragment size was assessed with a TapeStation and High Sensitivity D1000 ScreenTapes. The final fragment size of the libraries including adapters was on average 270 bp, due to the highly degraded nature of most samples. Up to 12 samples with similar library concentrations and fragment sizes were pooled for target enrichment. The dual-hybridisation procedure follows Hendriks et al. (2021) with both, the Angiosperms353 (Johnson et al., 2019, hereafter A353), and the Ochnaceae-specific (Schneider et al., 2020, hereafter OCHN) probe sets in the same reaction. To maintain equivalence between probes, a proportion of 3:1 *v/v* was used for each reaction, as A353 includes *ca.* 80,000 probes for 353 genes, while the OCHN set includes 19,398 probes for 275 genes (Schneider et al., 2020). Hybridisation was performed for 24h at 60*^◦^C*, followed by 12-14 (−18) cycles of PCR. Final products were analysed on a TapeStation to assess the fragment size, and calculate the final molarity. Sequencing was performed on an Illumina HiSeqX instrument at Macrogen (Europe, Netherlands) producing 2 *×* 150 bp paired-end reads. Raw data have been deposited in the European Nucleotide Archive (ENA, EMBL-EBI) under accession number PRJEB101313.

### 2.3 Data quality and assembly

Both sets of reads, new and previously-produced, were checked for initial quality with FastQC v.0.12.1 (Andrews et al., 2010). Reads were trimmed using Trimmomatic v.0.39 (Bolger et al., 2014) to remove adapter sequences and low-quality bases from the beginning and end of reads with a PHRED quality score *<* 20, using a 4-bp sliding window, and discarding reads trimmed *<* 30bp. An average of 9,897,591 reads per sample were kept after the trimming process for 96 samples sequenced in the present study. For sequences obtained from public repositories, an average of 1,759,526 (A353) and 3,857,892 (OCHN) reads were obtained for subsequent analyses (Table S3).

Mapping and assembling paired reads were performed using the pipeline HybPiper v2.1.3 (Johnson et al., 2016) including the following packages: BWA v.0.7.17-r1188 (Li, 2013) for mapping reads to the targets, Biopython v.1.81 (Cock et al., 2009) for handling reads, SAMtools v.1.17 (Danecek2021) for sorting reads, SPAdes v.3.15.5 (Prjibelski et al., 2020) for assembling reads into contigs, and Exonerate v.2.4.0 (Slater and Birney, 2005) for coding sequence extraction. Custom target files were constructed, one for A353 filtering the mega353 file (McLay et al., 2021) for Malpighiales, and one for OCHN based on the scaffold sequences used to design the probe set (see supplementary material for details). Custom target files resulted in higher capture success and longer sequences compared with the standard target file (Fig. S2; Baker et al., 2022). A total of 10 shared loci between both sets of probes were detected using the command-line version of BLASTn to avoid analysing the same regions twice (Supplementary material). Nine loci were removed from the A353 target file (the shortest targets), resulting in a final dataset of 618 nuclear genes, *i.e.,* 344 (A353) + 274 (OCHN).

Exon sequences for each sample and locus were retrieved using the HybPiper command retrieve sequences, which selects among contigs following three criteria: relative length compared to the target locus alignment, percent of identity between the coding sequence and the target, and depth of coverage. Sequences with about twice as much depth of coverage as the other contigs, and also with a higher percent identity were selected and identified by HybPiper with the suffix .main. Paralogous sequences (“paralogs”) were also retrieved using the HybPiper script paralog retriever.py, which generates one matrix per gene containing a single exon sequence for some samples (*i.e.,* the .main sequence), but multiple sequences for samples for which SPAdes detected more than one long-length contig (*≥* 75% of the reference sequence) that do not significantly differ from the .main sequence in coverage depth and percent identity (Johnson et al., 2016).

### 2.4 Filtering strategies

A total of eight datasets were analysed in order to assess the effect of missing data and paralogous sequences/genes on topological uncertainty and clade support for tribe Sauvagesieae. Loci and samples were selected for each dataset as follows (Table S2):

#### 2.4.1 Missing data

Under simulated conditions, filtering genes based on missing data does not impact or even reduce the accuracy of different methods to infer species trees (Molloy and Warnow, 2017). However, among empirical studies, there is no pattern regarding the effect of locus removal due to incomplete sampling (Hosner et al., 2016; Mclean et al., 2019; Sayyari et al., 2017). Nevertheless, alignments tend to be difficult when just few and not closely related species are sampled (Mirarab, 2019). Consequently, three filtered datasets were analysed here. First, all exons, which includes exon sequences for all samples and genes retrieved by HybPiper. Second, rm out, which includes exon sequences after removing partial sequences for each locus based on the assumption that very short contigs are more likely to be assembly artefacts, and are potentially difficult to align, even if they are genuine homologous sequences (Eiserhardt et al., 2022). Briefly, sequences shorter than the 25% of the median sequence length for that gene, or shorter than 150 bp, whichever was shorter, were removed. For very short loci (*<* 300 bp), sequences with at least 70 bp were kept. Sequences with more than 50% of unknown positions (N) and with no sequences higher than 150 bp between N positions were also removed (Fig. S3). Third, str mssdata, in which loci with less than 30% samples included, and samples with less than 30% of the total bp length were removed.

#### 2.4.2 Paralogs assessment

Multiple contigs per sample could be retrieved during SPAdes assembly for different reasons: allelic variation, the presence of different copies of a gene in a sample due to gene/genome duplication (*i.e.,* true paralogs) or hybridisation, but also through cross-contamination among libraries or sequencing errors (Morales-Briones et al., 2021a,b). Paralogous genes are prevalent in plants and constitute an important source of gene tree heterogeneity that has a negative impact on phylogenetic inference and support (Frost et al., 2024). Accordingly, we used two approaches to detect and analyse paralogs in our data. One implemented in HybPiper, which detects long-paralog and produces a fasta file containing all alternative contigs for each sample with at least 75% of the target sequence and high coverage depth (*i.e.,* exceeding 10 times competing contigs). Alternatively, HybPhaser v.2.1 (Nauheimer et al., 2021) uses an IUPAC ambiguity code to represent divergent nucleotides present in reads from all gene variants, and then quantifies those heterozygous sites (SNPs) per sample and gene. These two approaches are complementary because paralog detection in HybPiper is based on sequence length, depth and percentage identity among competing contigs, while HybPhaser quantifies heterozygous sites across all reads retrieved for a sample. Following those approaches, four additional datasets were analysed. First, all paralogs, which includes paralog sequences for all samples and genes retrieved by HybPiper. Second, inform paralogs, which includes exon sequences after removing samples from loci that were tagged as paralog by HybPiper, and for which all retrieved paralogs were not grouped in a clade. To choose those samples, initial gene trees were performed for each gene including all paralogs using the HybPiper script paralog retriever.py, followed by an automated inspection of the phylogenetic grouping of all paralogs for each sample. This test was implemented in the custom script removing paralogs.R (supplementary material). The .main sequence for samples for which all competing contigs (paralogs) formed a clade were kept, and remaining copies for that sample were removed for the specific gene (Fig. S4). This strategy avoids removing the entire gene with paralog warnings. Third, putative orthlgs, in which all loci tagged as paralogs by HybPiper were removed. Fourth, the dataset str mssdata prlgs, uses HybPhaser to identify and remove genes and samples with an unusual high proportion of heterozygous sites, *i.e.,* more than 1.5 times the interquartile range above the third quartile of the mean proportion of SNPs throughout the dataset, because they might have heterozygous sites due to the presence of paralogs or contamination (Nauheimer et al., 2021; Hendriks et al., 2023; Frost et al., 2024). Genes with more than one sample tagged by HybPiper as having paralogs were also removed from this dataset, keeping only loci for which there is no evidence of paralogy using both approaches.

Given the balance between loci, sample and site occupancy, clade support and topological differences among datasets, we generated an additional dataset (rmout infprlgs) combining two strategies: remove very short sequences such in dataset rm out, and remove paralogs using the grouping criterion such in the dataset inform paralogs.

### 2.5 Phylogenetic analyses

All datasets were aligned using the algorithm E-INS-i implemented in MAFFT v.7.520 (Nakamura et al., 2018) and trimmed to remove poorly aligned regions using TAPER v.1.0.0 (Zhang et al., 2021) with a cut-off value of -c 8 to perform a more conservative trimming. Gene tree inference was performed using IQ-TREE v.2.2.3 (Minh et al., 2020), with the best substitution model for each gene estimated by ModelFinder (Kalyaanamoorthy et al., 2017), using 100 different starting trees to explore the tree space, and a standard non-parametric bootstrap with 500 replicates to assess branch supports. Species trees for all datasets except all paralogs were estimated from the unrooted gene trees using Weighted ASTRAL-Hybrid v.1.15.1.3 (Zhang and Mirarab, 2022b) with 16 initial rounds of search, and a minimum threshold of 33% of bootstrap support. This method reduces the impact of quartets with low support and long terminal branches using a weighting scheme. Previous to the species tree estimation for all paralogs dataset, branches with low bootstrap support (*≤* 33%) in the gene trees were collapsed using the function nw ed of Newick Utilities v.1.6.0 (Junier and Zdobnov, 2010). The species tree for that dataset was inferred using ASTRAL-Pro2 v.1.15.1.3 (Zhang and Mirarab, 2022a) developed to use multi-copy gene trees as input. All paralogs for the same sample were forced to be grouped in the species tree by using a mapping file. Finally, all species trees were rooted in *Bonnetia stricta* + *Hypericum perforatum* using the function pxrr of the package Phyx (Brown et al., 2017).

The datasets were compared in terms of sample and site occupancy, median and variance of branch lengths, and median bootstrap support of the corresponding gene trees. Species trees were also compared using the quartet support (Q1) and the local posterior probability (LPP1) for the main topology obtained from ASTRAL, as well as the gene concordance factor (gCF) as the fraction of decisive gene trees concordant with each branch, the site concordance factor (sCF) as the fraction of decisive alignment sites supporting that branch, and the ultra-fast bootstrap support with 1000 replicates. These metrics were calculated in a concatenate-based analysis using IQ-TREE2 and an edge-linked partition model in which all partitions share the same set of branch lengths, but allowing each partition to have its own evolution rate in order to measure how each partition informs the topology. From the dataset rmout infprlgs, we also compared the metrics mentioned above but between probe sets (*i.e.,* A353 vs OCHN). Statistically significant differences between groups were assessed with the Kruskal-Wallis Test, followed by a Conover-Iman Test of Multiple Comparisons using the Bonferroni correction, implemented in the R packages stats (R Core Team, 2025) and conover.test (Dinno, 2024), respectively. Topological differences between species trees were measured by calculating the generalised Robinson-Foulds distances between trees, as implemented in the R package TreeDist v.2.9.1 (Smith et al., 2020). This method accounts for the information content of similar but not quite identical pairs of splits, which would contribute zero to tree similarity under the standard Robinson–Foulds measure.

### 2.6 Allelic divergence and locus heterozygosity to identify possible hybrid and polyploid lineages

After exploring gene trees from the dataset all paralogs we realised that in most of them, different sequences recovered for a given gene in many samples were grouped, instead of obtaining clades with all sequences for that gene and sample, as expected in the presence of alleles (Fig. S4). The high number of genes with paralog warning by HybPiper along with the above-mentioned pattern, suggest possible events of gene/genome duplication and hybridisation in Sauvagesieae. Then, we used HybPhaser to assess allelic variation in the tribe by calculating the average divergence between alleles (% allele divergence - AD), and the proportion of loci with divergent alleles (% locus heterozygosity - LH). These two metrics provide signals of hybridisation and duplication from target capture data. However, AD and LH values are relative to the dataset, sequencing quality, study group, and many other factors (Nauheimer et al., 2021). The expected range for ‘normal diploids’ is roughly *LH <* 90% and *AD <* 1%, while highly and recent polyploids tend to have high LH (*>* 80%) and AD (*>* 3.5%) values, and hybrids usually show high levels of LH (*>* 90%) and intermediate AD values (1-4%), that correspond to the divergence of the parental lineages (Hendriks et al., 2023). Establishing expected AD and LH values for old polyploids could be unreliable because duplicate genes are lost with time, decreasing LH values. AD for each sample was calculated as the relative SNP proportion across all genes (total bp), and LH was calculated as the percentage of genes with SNPs for each sample, after removing samples and genes with less than 10% of recovered target sequence length and proportion of samples recovered, respectively (Tables S3 and S4).

### 2.7 Phylogenetic networks to infer reticulate evolution in Sauvagesieae

To estimate probable hybridisation events within Sauvagesieae, a phylogenetic network was inferred using a pseudolikelihood approach in species networks applying quartets (SNaQ) (Solís-Lemus and Ané, 2016), implemented in the Julia package PhyloNetworks v0.16.4 (Solís-Lemus et al., 2017). Due to the computational limitations of SNaQ, we selected a subset of 28 species including those which showed signals of hybridisation according to HybPhaser (Table S4), and 592 loci from the dataset rmout infprlgs, after removing loci with less than 30% of species sampled. Network estimation was carried out with SNaQ using the species tree estimated with Weighted ASTRAL-Hybrid for the 28 species as starting tree, and considering the number of hybridisations to be h=0 to h=5. Each analysis included 20 independent runs to help optimisation explore more thoroughly the parameter space. We applied the heuristic criterion of gradient stabilisation to decide how many reticulations allow in the network (Solís-Lemus and Ané, 2016; Tiley et al., 2023). Finally, a bootstrap analysis was performed with 100 replicates, each running five parallel searches. Phylogenetic networks were plotted using the Julia package PhyloPlots.jl.

### 2.8 Estimation of ploidy level based on a site-based heterozygosity approach

To estimate the ploidy of Sauvagesieae species, we used the R package nQuack v.1.0.2 (Gaynor et al., 2024). This package implements an Expectation-Maximization algorithm using the normal, beta, and beta-binomial distributions fixing alpha, variance, or both parameters, to determine the most likely ploidy level of a sample (from diploid to hexaploid), given the empirical distribution of its allele frequencies. First, we used bwa-mem2 v.2.3 (Vasimuddin et al., 2019) to map trimmed reads for selected species (Table S1) to a reference sequence obtained from HybPiper for *Cespedesia spathulata*, one of the samples with the highest number of genes captured for both probe sets. Then, we converted the SAM files to sorted BAM files using SAMtools v.1.22.1 (Danecek et al., 2021). Before SNPs counting, poorly mapped reads and sites with more than 10% chance of being mapped to the wrong location were removed (-q 10). Given that the amount of variance or noise in the dataset affects the prediction accuracy, raw data was filtered by coverage using a minimum depth of 10, and by allele frequency using the nquire method implemented in the function denoise data, removing any site with frequencies below 0.15 or above 0.85. After filtering, 18 model types for all three distributions were tested, for a total of 54 models, followed by a model selection using the Bayesian Information Criterion (BIC) score, considering only diploid, triploid and tetraploid mixtures. To reduce the assignment error, models for higher ploidy levels were ignored (Gaynor et al., 2024; Corrêa dos Santos et al., 2017). The best model was used to estimate the ploidy level for each sample with 1000 bootstrap replicates using the function quackNboots. For samples with an absolute difference between the BIC score of the best model relative to the second-best model less than 10, bootstrap replicates were run using both competing models independently.

All analyses were performed using the UK Crop Diversity: Bioinformatics HPC Resource (Percival-Alwyn et al., 2025, https://www.cropdiversity.ac.uk).

## 3 Results

### 3.1 Library preparation, enrichment efficiency and gene recovery

By combining both universal (A353) and specific probe (OCHN) sets with a low hybridisation temperature, and using custom target files, we achieved 17% of reads on-target and 293*/*343 genes on average for A353, and 26% of reads on-target and 251*/*274 genes for OCHN across 96 samples (all of them corresponding from tribe Sauvagesieae). For A353, we recovered on average 179,734 bp and 386,121 bp for OCHN, representing 30% and 71% of the target length, respectively (Fig. 1). Herbarium samples showed comparatively lower DNA concentration, fragment size and capture efficiency (Fig. S1). Assembly performance was not affected by the initial gDNA concentration; thus it was possible to obtain hundreds of loci and base pairs from samples with as little as 10 ng of input DNA (Fig. S5), which is common in Sauvagesieae. In contrast, libraries with low concentration and small fragment sizes resulted in lower percentage of reads on-target and shorter sequences, even when the number of genes recovered was not affected by the library quality (Fig. S5). For only four species, we recovered less than 100 genes, all with less than 100,000 reads and a mean length captured of 21,443 bp (Table S3). Performance also differed between genera, with *Tyleria* showing low library concentrations, highly fragmented libraries, and the lowest number of genes recovered (409 genes and 263,404.5 bp on average; Fig. S1, Table S3). *Tyleria* is endemic to the Amazon-Guyana region and all samples included in this study came from old herbarium specimens, most of them probably fixed in alcohol. Although 15 samples showed less than 10% of the target length and genes recovered, most of them from old herbarium specimens; however, we successfully obtained hundreds of nuclear genes from pre-1950 samples, including types (Tables S1 and S3).

**Figure 1:**
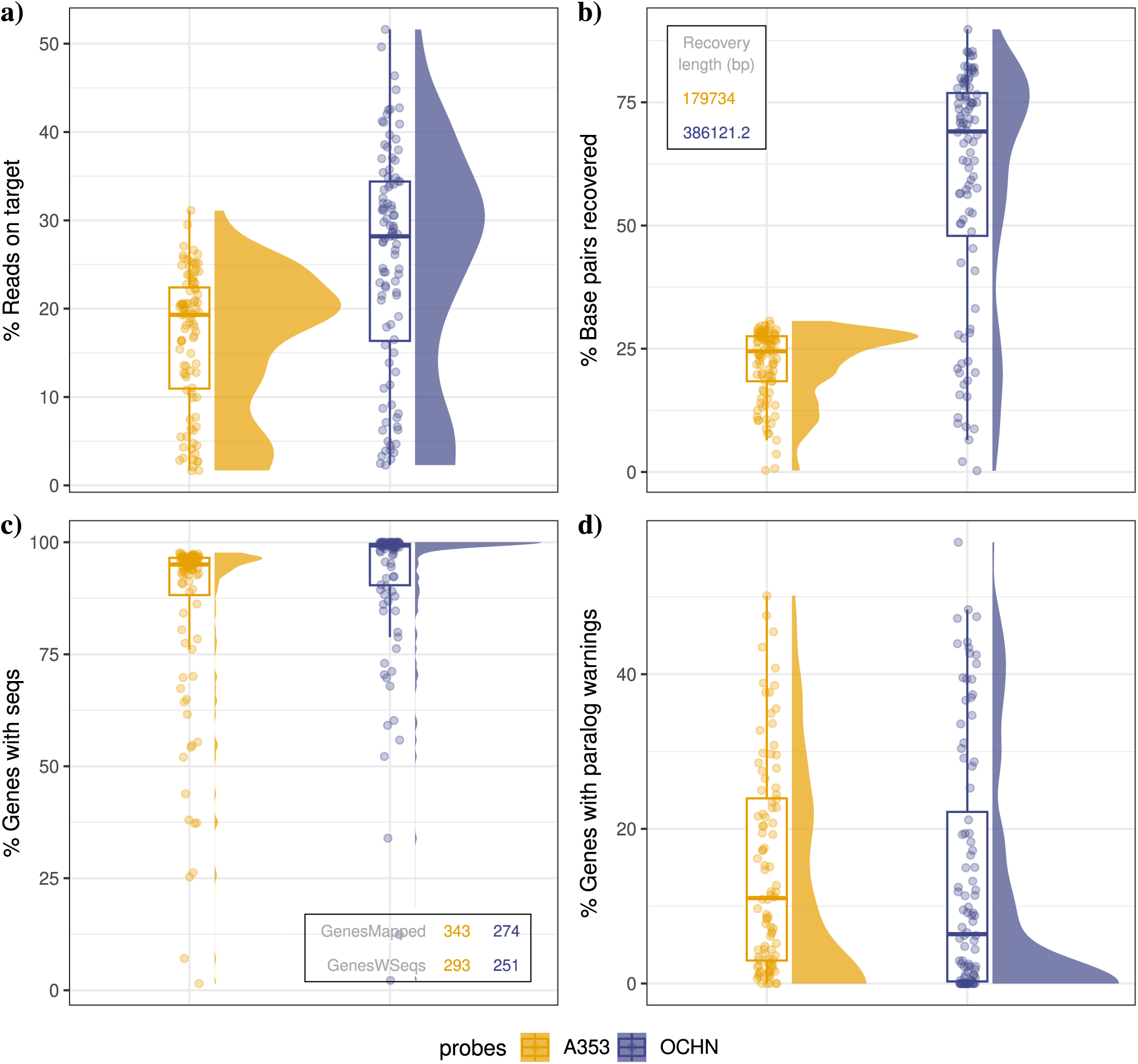
Capture and assembly summary using HybPiper. Differences between target-capture efficiency using the universal Angiosperms353 (A353) and the Ochnaceae specific probe set (OCHN) on 96 samples, corresponding to 73 species of tribe Sauvagesieae. a) Percentage of reads on target using the corresponding target files. b) Percentage of the total target sequence length recovered for each probe set. In the table, the mean sequence length recovered across samples per probe set. See Table S3 for details about target length for each gene. c) Percentage of genes mapped for which sequences were recovered. In the inside table, the mean number of genes mapped and the mean number of genes with sequence for each probe set across all samples are shown. d) Percentage of genes with sequences which had long and depth paralog warnings. The same pattern was found in the previously sequenced data using both probe sets separately (Fig. S6).

### 3.2 The effect of missing data and paralogs on gene-tree discordance

Filtering strategies drastically reduced the number of samples and sites per gene (*i.e.,* occupancy), as well as the number of genes informing the species trees, in particular for datasets putative orthlgs and str mssdata prlgs (Fig. 2a,b). The removal of extremely short sequences as in the dataset rm out (which avoids the loss of entire genes), and the removal of samples and genes with less than 30% sampling (dataset str mssdata) significantly increased the median branch support across gene trees, with a little effect on the median branch lengths (Fig. 2c,d). The median Q1 and LPP1 in the species trees, for both the overall tree and by clades, were not significantly different between datasets (Fig. 2e-i), except for the dataset all paralogs which showed the lowest median LPP1 and high variance across the nodes (Fig. 2f, i). However, gCF values showed no differences within clades, but statistically significant differences between clades (Fig. 2j), with *Cespedesia* clade, core *Tyleria* and *Sauvagesia* cladeA showing the lowest values regardless of the filtering strategy. The dataset str mssdata showed the lowest median Q1 among species trees, and for clades core *Tyleria* and *Sauvagesia* cladeA, but a slightly higher median gCF than the other datasets, suggesting an effect of filtering by missing data in those clades. Different levels of sample and gene removal due to the presence of missing data also resulted in slight differences across dataset topologies, mainly in the shallow nodes (Figs. S7-S17).

**Figure 2:**
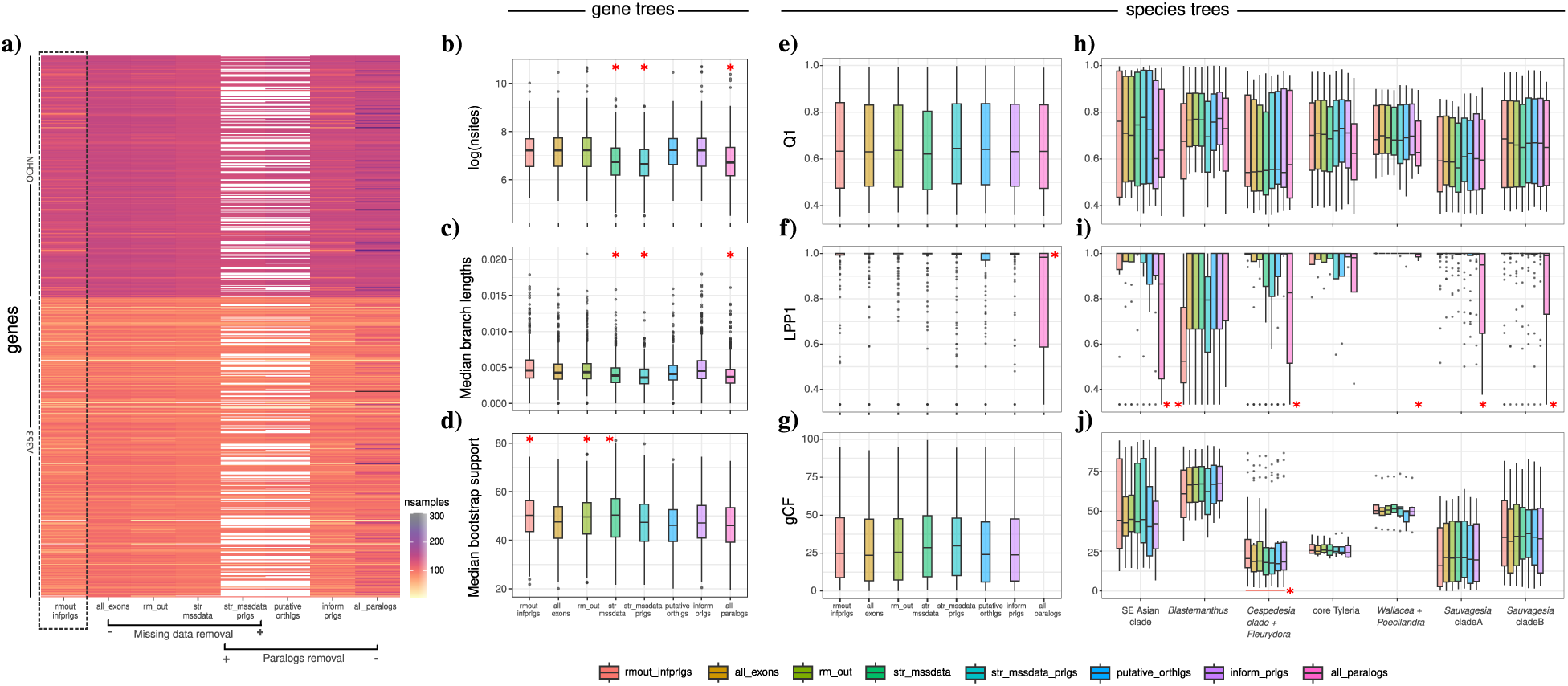
Comparison of gene-tree summary metrics from IQ-TREE2 (a-d), and branch support metrics for the species trees obtained using ASTRAL (e-j), for the eight datasets. a) Sample occupancy per gene (rows) in each dataset; empty rows represent lost genes. b) Median number of sites across genes in each dataset measured from the aligned matrix for each gene. c) Median gene-tree branch lengths across genes for each dataset. d) Median gene-tree bootstrap support across gene trees for each dataset. e-g) Median values for all nodes in the species trees for each dataset. h-j) Median values summarised by main clades in panels. Plotted variables correspond to: quartet support for the main topology (Q1) and local posterior probability for the main topology (LPP1) obtained using ASTRAL, and gene concordance factor (gCF) calculated using IQ-TREE2. The dataset all paralogs have no values for gCF because this support metric is calculated on a tree produced by maximum likelihood on a concatenated alignment, and different copies of the same sample in a gene do not correspond with copies with the same name in other genes (*e.g.,* SPRL 195.0; see Fig. S4). Red asterisk indicates statistically significant differences between groups (*p <* 0.05) according to a Kruskal-Wallis Test followed by a Conover-Iman Test of Multiple Comparisons Using Rank Sums.

Although the presence of missing data did not significantly affect both topology and support of gene and species trees, filtering strategies used to deal with paralogs resulted in notable differences among species trees in shallow and deep nodes (Figs. S7-S17). The dataset all paralogs resulted in the most dissimilar species tree topology among datasets (Fig. S8), and showed the lowest LPP1 values, for the overall tree and by clades (Fig. 2i). In addition, gene trees for this dataset had a low median branch support and number of informative sites (Table S2; Fig. 2b), suggesting lower quality alignments. Removing paralogous sequences instead of removing entire loci, as in the datasets rmout infprlgs and inform prlgs, resulted in higher values of Q1 and gCF for SE Asian clade, *Cespedesia* clade and *Sauvagesia* cladeB, whereas removing all loci with paralog warnings as in the dataset putative orthlgs neither significantly increased the branch support nor decreased the topological discordance (Fig. 2c,d). The differences in median Q1 values among datasets found for *Blastemanthus* clade suggest an increase in conflict between gene trees when both missing data and paralogs are filtered at the same time, probably due to the original low amount of data for this clade (Fig. 2h-j; Table S3).

A high overlap between paralogs identified by HybPiper and HybPhaser was observed, with 174 genes for A353 and 124 for OCHN reported by HybPiper as having more than one contig with high coverage, length and percentage of identity, and 103 (A353) and 176 (OCHN) with high mean proportion of SNPs (*>* 2%) across all samples according to HybPhaser (Table S5). However, 30% of genes with paralog warnings by HybPiper showed a low proportion of SNPs (SNPs *<* 2%), suggesting the presence of probable alleles instead of true paralogs for some samples in those genes. We found evidence for this hypothesis after examining the phylogenetic grouping of competing contigs for 18 of those genes (Table S5). In contrast, HybPhaser detected at least 91 genes based on SNPs proportion which have no competing contigs (potential paralogs) detected by HybPiper (Table S5). The level of overlap between genes/species tagged as paralogs by both approaches depends on the selected thresholds, as expected. When the threshold is the outlier percentage of SNPs across samples, just 21 genes are removed by HybPhaser, but using 2% SNPs as suggested by Hendriks et al. (2023) as a high level of SNP proportion, 271 genes are recognised as paralogs. However, the overall number of SNPs in a dataset depends on the group, the sequencing quality, and random error. Consequently, using the outlier threshold seems to be more objective, despite the fact that it can underestimate the number of paralogs.

The removal of paralogs detected using both approaches (dataset str mssdata prlgs) led to the loss of 276 loci (44% of total loci) and a drastic reduction in the number of samples and sites per gene (Fig. 2a-b). This stringent filtering did not significantly improve the branch support of the gene trees nor the species trees (Q1, LPP1 and gCF) for most clades (Fig. 2b-j). The dataset rmout infprlgs resulted in the best balance between data loss, median branch support among gene trees, and higher Q1, LPP1 and gCF values for the species tree (Fig. 2), while removing putative paralogous sequences for which it is not possible to properly establish their homology. For that reason, further analyses were performed using this dataset.

### 3.3 Contribution of each probe set to the species tree

Although the OCHN probe set (275) includes fewer loci than A353 (353), OCHN loci are longer and resulted in gene trees with higher mean bootstrap support (Fig. 3a). This pattern was consistent across all filtering strategies (Fig. S18). However, the increase in gene-tree support was not directly reflected in the Q1 and LPP1 values of the resulting species trees (Fig. 3b-d). Topologies inferred using each probe set independently were also broadly congruent and recovered the major clades within the tribe, but showed some discordant relationships mainly within the *Sauvagesia* cladeA and the *Cespedesia* clade (Fig. S19). Although A353 loci produced a species tree with lower gene concordance factor (gCF) and slightly lower Q1 support, the integration of A353 and OCHN loci improved both locus and taxon sampling, and increased the resolution of some nodes within the *Sauvagesia* cladeB, *Rhytidanthera*, and the clade uniting *Sauvagesia elegantissima*, *S. bryoclada* and *S. spicata* in the *Sauvagesia* cladeA (Figs. S19-S21).

**Figure 3:**
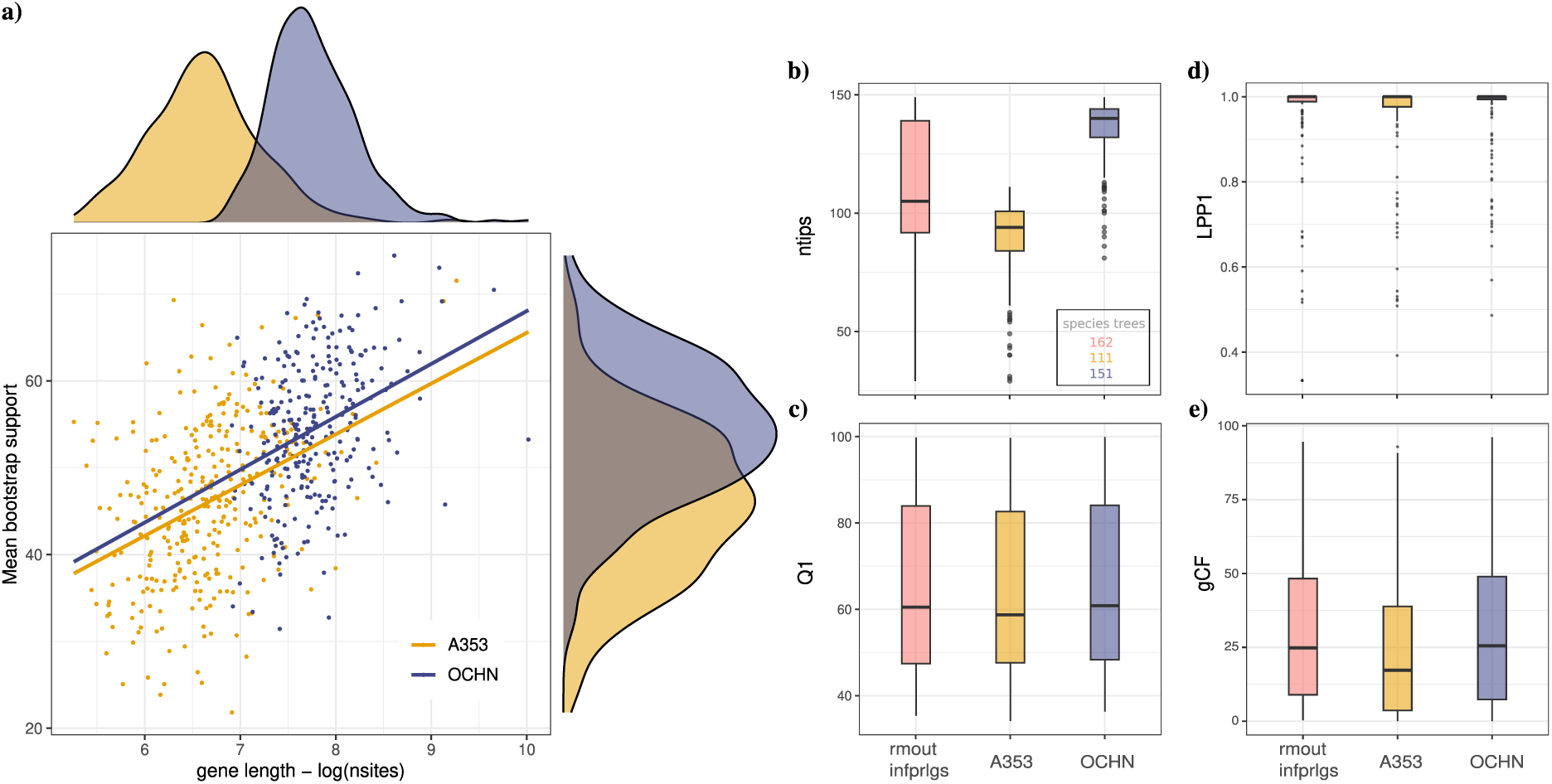
Comparison of gene-tree summary metrics from IQ-TREE2 (a,b), and branch support metrics for the species trees obtained using ASTRAL (c-e), for the dataset rmout infprlgs but analysing loci from the two probe sets independently. a) Gene length in number of sites for each locus alignment vs the median bootstrap support for the corresponding gene tree. The same pattern was observed across filtering strategies (Figure S18). b) Median number of tips across gene trees for the complete dataset rmout infprlgs and by probe set. In the table, the total number of tips in the corresponding species tree. c-e) quartet support for the main topology (Q1) and local posterior probability for the main topology (LPP1) obtained using ASTRAL, and gene concordance factor (gCF) calculated using IQ-TREE2 for the complete dataset rmout infprlgs and by probe set.

### 3.4 Phylogenetic relationships

The backbone of the species tree reconstructed using ASTRAL was consistent across datasets (Fig. S7). Regardless of the low quality of some samples, especially those from old herbarium specimens, and the overall high gene-tree discordance, all analysed species trees retrieved the major clades previously reported for Sauvagesieae (Schneider et al., 2014, 2020, 2021; Reinales and Parra-O., 2020) (Fig. 4): The *Cespedesia* clade which is sister to *Fleurydora* and both to *Blastemanthus*, the small clade comprising *Poecilandra* and *Wallacea*, the SE Asian clade, and the two clades in which *Sauvagesia* species were segregated. *Sauvagesia* cladeA including all species originally described in *Lavradia*, and the *Sauvagesia* cladeB sister to *Adenarake*. However, the internal branches of both *Sauvagesia* clades showed a strong tree discordance across datasets and among gene trees, as suggested for the low Q1 values and the high differences between gCF and sCF, mainly in the deeper nodes (Figs. 4 and S17), which was slightly reduced after removing missing data and paralogs (Fig. 2).

The most remarkable difference among datasets, and compared to previous phylogenetic studies was the position of *Tyleria*. This genus was recovered as polyphyletic in all datasets, even after different levels of paralog removal and a moderate removal of missing data (Figs. S7-S17). All samples sequenced by Schneider et al. (2020) and one sequenced in this work were grouped as a clade (core *Tyleria*), but the position of this clade is also unclear with two alternatives: core *Tyleria* sister to the SE Asian clade, or sister to the clade including *Sauvagesia* and *Adenarake*. Two datasets support the second alternative, str mssdata prlgs in which missing data and paralogs were removed, and all paralogs which includes all competing contigs in the same alignment. However, this position is weakly supported (Q1=45, LPP1=0.9, gCF=0.3; Figs. S7, S13, S15 and S17).

Within the *Cespedesia* clade, *Rhytidanthera* was found to be sister to a clade formed by *Godoya*, *Krukoviella* and *Cespedesia* with low tree discordance and moderate support (Q1=86, LPP1=1, gCF=24.8, sCF=47). However, there is high discordance in the relationships between *Godoya* and *Krukoviella* because alternative topologies have almost the same frequency that the main one (Fig. 4). All samples of *R. regalis*, including the isotype *Idrobo & Schultes871*, formed a clade with moderate discordance and about 35% of the informative sites for this branch supporting its status as a valid species (Q1=52, LPP1=1, gCF=3.5, sCF=35.5). The relationships among the remaining samples of the genus were not resolved because the two samples of *R. sulcata* did not form a clade and *R. splendida* was recovered as paraphyletic, but with strong tree discordance and low support given that the gCFs were less than half of the sCFs, and Q1 values were less than 50 in most nodes (Figs. 4 and S8). Due to the good sampling achieved for these samples, the high tree discordance is probably caused by reticulation events. *Godoya antioquiensis* was recovered separately from the clade formed by *G. obovata* samples with high support (Q1=71, LPP1=1, gCF=48, sCF=43.1).

Within the SE Asian clade, *Schuurmansia* + *Schuurmansiella* are sister to *Euthemis*, which is sister to a clade formed by *Neckia* and *Indovethia* + *Indosinia* with high support despite gene tree discordance (Q1=98, LPP1=1, gCF=45.7, sCF=46.6; Fig. 4). Those relationships are also consistent across datasets (Figs. S7-S17). However, the position of *Neckia* in the species tree is weakly supported by the gene trees (Q1=42; Fig. 4), with an alternative position as sister to *Euthemis* and the clade formed by *Indovethia* + *Indosinia*, which is supported by about 30% of both genes, and informative sites (Fig. S17).

Within the *Sauvagesia* clades, we also observed strong gene tree discordance, regardless of the different level of missing data and paralog removal (Fig. S9-17). However, some noteworthy clades were recovered in all datasets with low conflict (Fig. 4): samples of *Sauvagesia glandulosa* were separated from samples of *S. rubra* (Q1=82, LPP1=1, gCF=43.7, sCF=46.5), recently suggested as different species by Queiroz-Lima et al. (2023); samples for *S. vellozii*, the type species of the genus *Lavradia* (synonym of *Sauvagesia*), were grouped (Q1=79, LPP1=1, gCF=39.6, sCF=45.4) and recovered as sister to *S. capillaris* within the *Sauvagesia* cladeA (Q1=57, LPP1=1, gCF=10.9, sCF=31.1); samples of *S. linearifolia* from Colombia are sister to *S. deflexifolia* (Q1=87, LPP1=1, gCF=62.7, sCF=27.8), and were separated from the Brazilian specimens, which are sister to *S. lanceolata* (Q1=87, LPP1=1, gCF=55.5, sCF=41.7); samples of *S. erecta* var. *coriacea* formed a clade sister to *S. racemosa* (Q1=77, LPP1=1, gCF=35.5, sCF=44.1), and were separated from other samples of *S. erecta*. In addition, *Sauvagesia africana*, sequenced for the first time, was retrieved as sister to *Sauvagesia* cladeB and *Adenarake* in all datasets, with moderated support (Q1=49-56, LPP1=1, gCF=5.7, sCF=42.7; Figs. S9-17).

For few species across the main clades, different samples analysed for the same species were not retrieved as a clade such as *Schuurmansia henningsii* and *Neckia serrata* within the SE Asian clade, *Sauvagesia amoena* in *Sauvagesia* cladeB, *Sauvagesia spicata* and *Sauvagesia ribeiroi* within *Sauvagesia* cladeA, *Poecilandra pumila*, and *Rhytidanthera sulcata* in the *Cespedesia* clade, but in all cases with high tree discordance and low support (Fig. 4).

### 3.5 Allelic divergence and signals of polyploidisation and hybridisation in Sauvagesieae

After assessing allelic variation by calculating the proportion of loci with divergent alleles (LH), and the average divergence between alleles (AD) on 608 nuclear loci using HybPhaser, most samples for the SE Asian clade showed the expected range for diploid species, with roughly *LH <* 90% and *AD <* 1% according to Hendriks et al. (2023). However, a significant number of samples showed signs of recent polyploidisation, especially within *Sauvagesia* cladeA, *Cespedesia* clade, *Sauvagesia* cladeB, and core *Tyleria*, which have high LH (*>* 75%) and AD (*>* 2%) values (Fig. 6a and Table S4). About 18% of the samples showed signs of hybridisation, mainly within the genera *Rhytidanthera*, *Sauvagesia* and *Tyleria*, with very high values of LH (*>* 90%) and intermediate values of AD (1.5-3.5%). Data from both probe sets showed a similar pattern of allelic divergence (Fig. S22). However, A353 genes have lower values of AD and LH, *i.e.,* lower percentage of SNPs across genes and samples, respectively.

**Figure 5:**
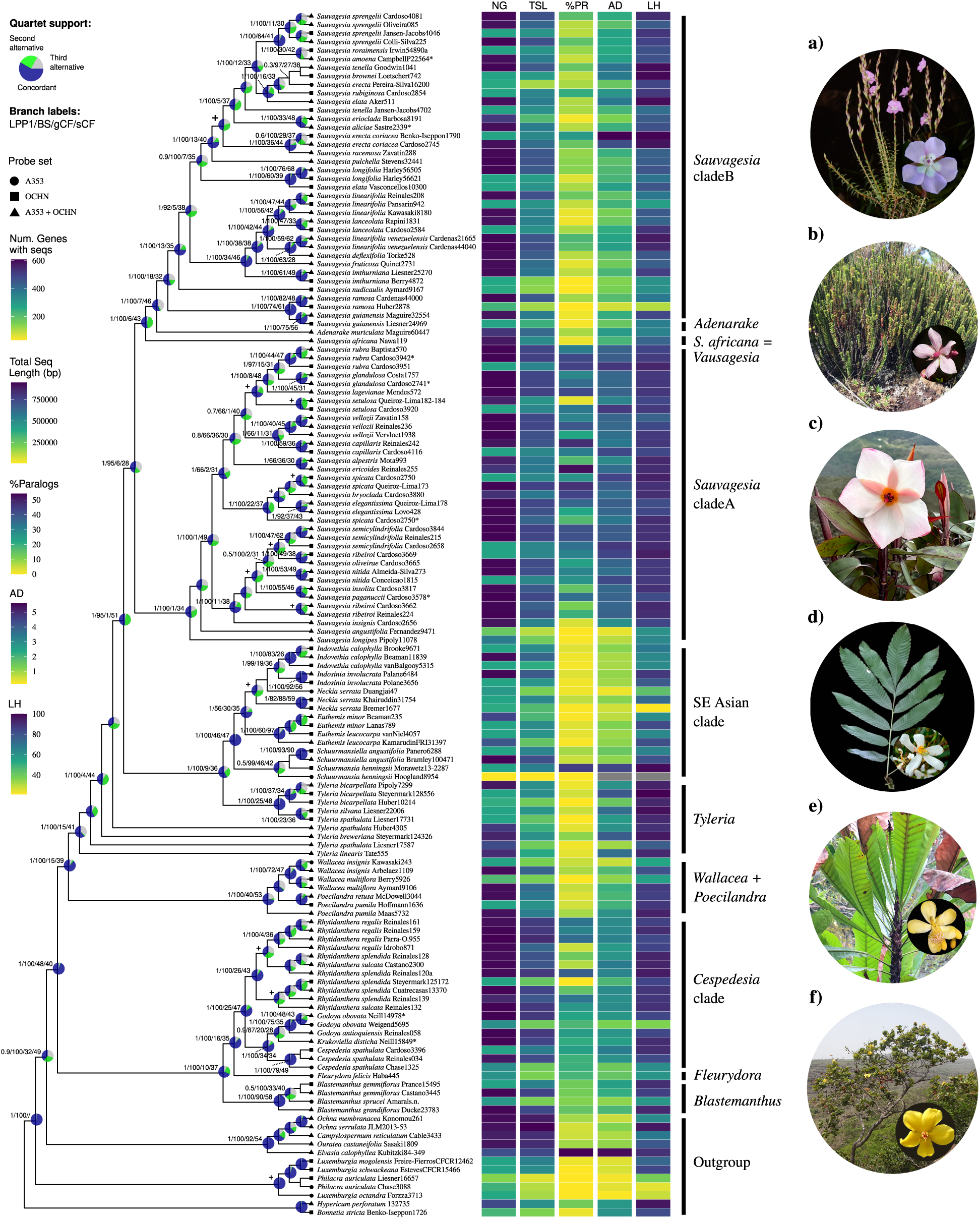
Species tree for Sauvagesieae resulted from summarising 608 gene trees using Weighted ASTRAL-Hybrid on exon sequences retrieved by HybPiper after removing extremely short sequences and paralogs (dataset rmout infprlgs). Pie charts indicate the degree of discordance between gene tree as quartet support. The proportion of input gene trees satisfied by the species tree Q1 (blue), and alternative frequent topologies Q2 and Q3 (green and grey respectively). A similar proportion of Q2 and Q3 suggests ILS, while different proportions of the two alternative quartets indicates probable hybridisation or polyploidisation events. Branch support is reported as local posterior probability for the main species tree from ASTRAL, as well as ultrafast bootstrap (BS), and gene (gCF) and site (sCF) concordance factors from IQ-TREE2. Colour bars represent the number of genes (NG), total sequence length (TSL), and the percentage of genes with paralog warnings (PR) recovered by HybPiper for each sample, and the allele divergence (AD) and locus heterozygosity (LH) percentage per sample calculated using HybPhaser. Black crosses represent conflicting nodes between ASTRAL tree and concatenated-matrix tree (Fig. S17) inferred to calculate the concordance factors. a) *Sauvagesia deflexifolia Figueira1805*. b) *S. ericoides* Reinales255 (Flower detail from Cardoso4113). c) *Tyleria tremuloidea* Barbosa-Silva1978. d) *Rhitidanthera sulcata* Reinales132. e) *Cespedesia spathulata* Ballen550. f) *Fleurydora felicis* from RBGK-POWO.

**Figure 6:**
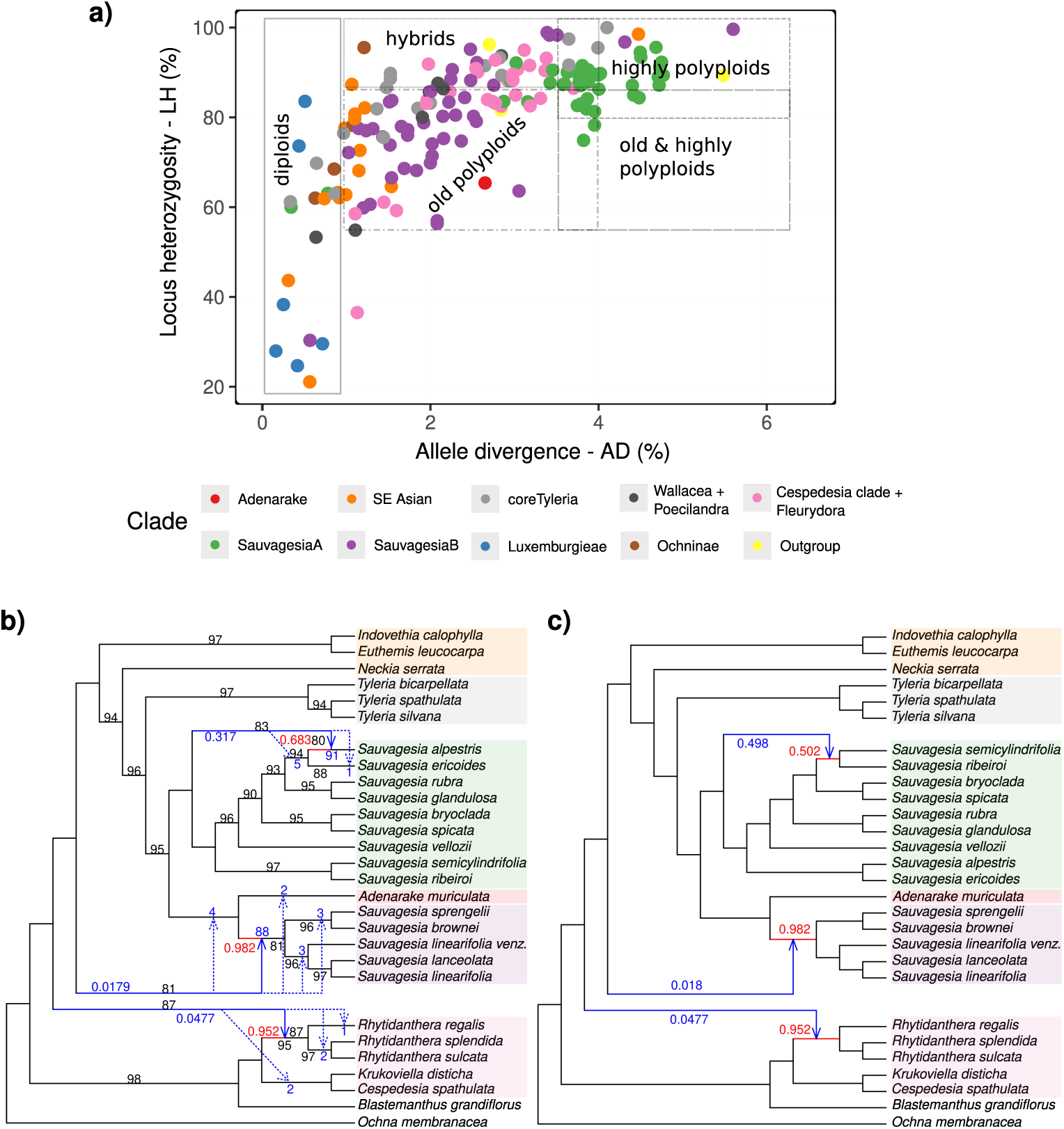
Allelic variation and phylogenetic networks showing signs of polyploidisation and reticulation in Sauvagesieae. a) Scatterplot of the allele divergence and locus heterozygosity for all samples calculated using HybPhaser. Samples and genes with less than 10% of sampling, and samples with an outlier SNP proportion for each gene were removed. ‘Normal diploids’ show roughly *LH <* 90% and *AD <* 1%, while highly and recent polyploids tend to have high AD (*>* 3.5%) and LH (*>* 80%) values, and hybrids usually show high levels of LH (*>* 90%) and intermediate AD values (1-4%). AD and LH values are relative and can vary depending on the dataset, sequencing quality, study group, and many other factors. b-c) Phylogenetic networks inferred using SNaQ. b) Best network with *Hmax* = 3. Numbers on branches represent bootstrap values lower than 100; for the main topology (black), for recipient edges (blue). Any alternative recipient positions for a reticulation event is shown as a dotted line. c) Alternative network from the network file which contains multiple candidate networks resulted from perturbations of the best network. It shows a different position of the major hybrid edge within *Sauvagesia* cladeA. In both cases, inheritance proportion (*γ*) values are presented for the optimal network; *γ* for the major edges (red) and minor edges (blue), depicting their point estimates.

### 3.6 Phylogenetic networks

The phylogenetic network analysis performed on a subset of 592 loci and 28 species (representing the main clades of Sauvagesieae) suggested three events of reticulation within the tribe (Fig. 6b). Gradient stabilisation of pseudodeviance scores for runs *h*0 to *h*5 showed the inflection point at the run with three hybridisation events, and consequently was selected as the best network (Fig. S23). The most frequent reticulation event has its origin in the ancestral lineage of the *Sauvagesia* cladeA (BS=83, inheritance proportion of *γ* = 0.317), to *Sauvagesia alpestris*, the most probable recipient (BS=91, *γ* = 0.683). The second most frequent minor edge has its origin in an extinct or unsampled taxa, in the branch of the clade uniting *Sauvagesia*, *Tyleria* and the SE Asian clade, and the recipient lineage in the branch of *Rhytidanthera* (BS=95), with a high inheritance proportion (*γ* = 0.952). Finally, the less frequent minor edge also originated in an unsampled or extinct ancestral branch, to *Sauvagesia* cladeB (BS=88, *γ* = 0.982). The main source of uncertainty in the second and third hybridisation events probably lies with the recipient branches, with more than four alternative recipients in each case, most of them showing low bootstrap values (Fig. 6b). In addition, these reticulations had extremely low gamma values for the donor, and probably represent optimisation errors (C. Solís-Lemus, *pers. comm.*).

Although the best network was chosen because of its pseudodeviance value, SNaQ also does a round of topological perturbations on the best network to help check whether other candidate networks are found that way. Among these networks, we observed the presence of a network that points to a reticulation event inserted onto the branch leading to *Sauvagesia semicylindrifolia* and *Sauvagesia ribeiroi* (Fig. 6c), nonetheless having a higher pseudodeviance than the one recovered as point estimate of network topology, with realistic inheritance proportions for both donor and recipient.

### 3.7 Ploidy estimation

After removing highly variable (noise) and extreme allele frequencies determined based on the inferred alpha parameter and allele frequency histograms, at least 11*/*80 samples resulted in low median coverage per site (*<* 50), and consequently it was not possible to accurately estimate their ploidy (Table S6). For the remaining 69 samples, no single model was selected as the best for all samples, being the normal distribution with only alpha and variance free, and a uniform mixture, the most common (normal-uniform fixed 2), followed by the model with the same distribution but only alpha free (Table S6). Most of the samples were characterised as diploids with high bootstrap support (*>* 85%) across main Sauvagesieae clades, and 13 samples were suggested as tetraploids, but with low to moderate support (25-75%). At least 10 samples showed a high uncertainty between diploid and tetraploid ploidy, including members of *Cespedesia* clade such as *Rhytidanthera sulcata*, *Fleurydora felicis*, *Sauvagesia alpestris*, *Sauvagesia capillaris* and *Sauvagesia glandulosa* of *Sauvagesia* cladeA, *Sauvagesia linearifolia* from Colombia, *Sauvagesia sprengelii* within *Sauvagesia* cladeB, *Neckia serrata*, and most of *Tyleria* species; some of them even have a high per-site coverage (Table S6).

## 4 Discussion

### 4.1 Dual-hybridisation, assembly, and paralogy assessment

A variety of genome-scale methods, such as target capture, have been developed to optimise the balance between the amount of data produced and sequencing costs, while increasing phylogenetic resolution and improving branch length estimation in studies across a wide range of taxonomic levels (McKain et al., 2018; Weitemier et al., 2014; Johnson et al., 2019). However, the larger the amount of data obtained (reads and samples), the more challenges the researchers have to face in data processing using tools that are generally conceptualised for diploid organisms and single-copy loci (Tiley et al., 2024; Morales-Briones et al., 2021b). Events of gene duplication and hybridisation are common in plants, and usually appear in target capture data as tree discordance either between genome topologies (*i.e.,* plastid vs. nuclear) or among gene trees (Cai et al., 2021; Morales-Briones et al., 2021b). Then, obtaining hundreds of independent loci allows us to test the presence of those phenomena in the group of interest that otherwise would be overlooked.

Combining universal and lineage-specific probes in a dual-hybridisation reaction allowed joining sampling and sequencing efforts to infer a densely sampled phylogenetic tree for Sauvagesieae, including all genera, and *ca.* 90% of the tribe’s extant species richness, without requiring deeper sequencing than one might have applied for a single probe set. Using this strategy, we achieved a higher percentage of reads on-target (22%) and a higher number of captured loci (450) on average per sample compared to those obtained by single hybridisations as in Schneider et al. (2020) which used the OCHN set (2-9.6% reads and 79-222 loci depending on the dataset), and Shah et al. (2021) which used the A353 and OCHN probes independently (11.1% and 256 loci for A353, and 47% and 241 loci for OCHN). This result reinforces that dual-hybridisation is a good alternative to optimise laboratory work and sequencing costs while increasing data output. However, reducing the hybridisation temperature to 60*^◦^C* to improve the capture sensitivity for both probe sets could allow partial matching between probes and targets, increasing the chance of capturing paralogous sequences (Forthman et al., 2023; Cruz-Dávalos et al., 2017). Nonetheless, the high sequencing depth used in this study increased coverage, reducing the proportion of stitched contigs, and consequently controlling the number of paralogs (Frost et al., 2024). The use of paired-end reads also improved confidence in the accuracy of variant calling, reducing the noise introduced by sequencing errors (Kotlarz et al., 2024).

The number of reads obtained per sample is correlated with the total sequence length captured, as expected, but also with the percentage of paralogs retrieved by HybPiper. This pattern was consistent for both probe sets (Fig. S24) independently, suggesting that more data allow the uncovering of paralogs that go unnoticed in samples with fewer data. Paralogs are more frequent when the taxa under study are distantly related to those used in the design of the probes (Moore-Pollard et al., 2024), as is the case in the present study, even with the specific probe set, which did not include any Sauvagesieae species (Schneider et al., 2020). In addition, when the read depth is high, reads of additional gene copies may pass the minimum read depth filter during read assembly, resulting in multiple contigs per locus (Ufimov et al., 2022).

A detailed analysis of paralogs is desirable in most cases, even more so when we expect the presence of polyploidisation or reticulation events that can result in gene alignments containing contigs from different subgenomes. Removing entire loci due to the presence of paralogs did not show a significant effect on reducing GTEE (Fig. 2). That strategy is unsuitable for groups with recent WGD events, as deleting loci with paralog warnings will cause substantial data loss (Ufimov et al., 2022). Pipelines such as HybPiper are not designed to distinguish between allelic variation and true paralogs, because during the assembly process only the most frequent nucleotide sequence is considered and alternatives are rejected (Prjibelski et al., 2020; Morales-Briones et al., 2021b). As a result, variable positions are collapsed into a single base call, losing information related to heterozygosity (Tiley et al., 2024; Hendriks et al., 2023). Consequently, analysing together the paralog warnings by HybPiper, allelic variation by HybPhaser, and paralogs all clade test allowed us to identify probable alleles, avoiding discarding valuable information (Table S5).

The removal of missing data and paralogs did not significantly improve the clade support nor reduce the tree discordance measured as the gene concordance factor (gCF) and the quartet frequency (Q1), respectively (Figs. 2 and 4). Instead, removing samples and entire loci because paralog warnings could increase the chance of recovering erroneous branches (GTEE), as observed in the datasets str mssdata prlgs and putative orthlgs, which have the lowest median branch support among gene trees (Fig. 2d). The risk is higher when those genes have high occupancy and increase the quartet scores for either of the alternative species trees (Gatesy et al., 2019). The reason is that in the presence of uneven taxon occupancy, the resulting ASTRAL trees may be driven by a handful of genes, and could change if only a couple of genes with the above-mentioned conditions are removed from the dataset (Mirarab, 2019; Ufimov et al., 2022). Consequently, a probable gain in gene tree quality by removing partially sampled genes, and entire genes for paralog warnings was not enough to compensate for data reduction (Molloy and Warnow, 2017; Morales-Briones et al., 2021b). However, very short sequences are more likely to be assembly artefacts and therefore could increase species tree error (Sayyari et al., 2017; Hosner et al., 2016). We showed that filtering fragmentary sequences but keeping as many genes as possible as in the dataset rmout infprlgs, improved gene tree estimation, and slightly reduced the tree discordance at least for some clades. The all exons dataset resulted in similar values of support and low gene tree conflict compared to rmout infprlgs, but there is no argument to choose among competing contigs in favour of the ‘.main’ sequence. In addition, the preferential selection of the longest contig with the highest coverage during assembly further contributes to underestimate the presence of paralogs and increases chimeric assembly (Frost et al., 2024). Thus, removing ‘non-monophyletic’ contigs for each gene turned out to be the best informed strategy. Combining this strategy with a quartet-based method for species tree reconstruction such as wASTRAL, mitigates the impact of GTEE on the species tree by weighting gene trees based on their branch lengths and support values (Zhang and Mirarab, 2022b).

Currently, numerous empirical studies have employed different approaches to detect and deal with paralogs using target-capture data (*e.g.,* Frost et al., 2024; Morales-Briones et al., 2021a; Hendriks et al., 2023). Their effect in reducing GTEE depends on the quality of the dataset and the complexity of the molecular evolution of the group under study. However, the growing availability of long-read sequencing data substantially improves paralog detection by allowing the accurate recovery of multiple gene copies and the assembly of haplotype-resolved genomes (Tiley et al., 2024; Ning et al., 2024).

### 4.2 Combining evidence of allele divergence, locus heterozygosity, allele frequencies, and quartet frequencies to hypothesise reticulation and recent polyploidisation in Sauvagesieae

The HybPhaser analysis suggested potential hybrids and polyploids, mainly within *Sauvagesia* cladeA and *Cespedesia* clade (Fig. 6a, Table S4), based on the quantification of the proportion of SNPs across all genes (allele divergence - AD), and the percentage of genes with SNPs (locus heterozygosity - LH). Both lineages are endemic to the mountain regions of South America, *i.e.*, the northern Andes and the Brazilian Espinhaço Range (Reinales and Parra-O., 2020; Queiroz-Lima et al., 2023). This result is in line with the high number of genes with more than one copy identified by HybPiper (Fig. 4). In contrast, tree taxa inhabiting the tropical forest, such as *Wallacea* from Amazon and *Euthemis* from SE Asia were suggested as normal diploids. nQuack also suggested that most of Sauvagesieae species are currently diploids (Table S6), including all members of *Cespedesia* clade, in contrast to their high allele divergence and locus heterozygosity. Although heterozygosity tends to be different (usually higher; Soltis and Soltis, 2000) in polyploids compared to diploids, predicted tetraploids could show lower percentages of polymorphic sites compared to related diploids (*e.g.*, *Dioscorea*, *Arabidopsis*; Viruel et al., 2019; Paape et al., 2018) due to strong selection on key genes, niche specialisation and smaller effective population sizes, among other mechanisms (Wright et al., 2015). Hence, signs of polyploidisation from HybPhaser based on the percentage of polymorphic sites, and from nQuack based on the allele frequency distribution should be carefully compared.

Most topological incongruences among filtering strategies also involve species with a high number of paralogs that were suggested as potential hybrids due to high AD and LH values, such as *Sauvagesia ribeiroi*, *Sauvagesia spicata*, and *Sauvagesia alpestris* (Figs. 4 and 6; Table S4). However, other samples with an extremely high amount of paralogs but without topological conflicts such as *Sauvagesia glandulosa* and *Sauvagesia rubra*, could represent highly polyploid species instead of hybrids, as suggested by their high AD but low to intermediate LH (Fig. 4; Table S4). This hypothesis was supported by the phylogenetic network analysis in which a probable reticulation event was inferred with the major hybrid edge in *Sauvagesia alpestris* (Fig. 6b), and other in the clade *Sauvagesia ribeiroi* + *Sauvagesia semicylindrifolia* (Fig. 6c), but no signal of reticulation for the *Sauvagesia glandulosa* clade (Fig. 6). None of these species were predicted as polyploid by nQuack (Table S6), except *Sauvagesia alpestris* which showed a high uncertainty between tetraploid and diploid, even despite its high per-site coverage. However, these clades could have undergone a diploidisation process, which is common after allopolyploidisation events resulting from hybridisation in angiosperms (Leitch and Bennett, 2004; Cai et al., 2019). Furthermore, the sequences produced in this study correspond to genes of low to single copy, which are prone to rapid diploidisation following WGD events (De Smet et al., 2013). Allele phasing can improve the power of network methods such as SNaQ by enabling the distinction between hybridisation and polyploidisation processes, and allowing the detection of single reticulation events in polyploids, even when only one parental lineage was sampled (Tiley et al., 2024).

Hybridisation and polyploidisation have been suggested as important mechanisms driving plant diversification through increasing genomic diversity, creating morphological novelties, and enhancing evolutionary success during colonisation of new environmental niches (Fawcett et al., 2009; Abbott et al., 2013; Hegarty and Hiscock, 2008; te Beest et al., 2011; Soltis, 2013). The period of strong climatic instability during the last 5 Ma (Pliocene), when most of the *in situ* speciation events occurred in the Espinhaço Range lineages (Vasconcelos et al., 2020), provides a plausible scenario to explain how secondary contact between species allowed gene flow and hybridisation during the coldest periods, followed by a period of isolation in which diversity was constrained by environmental factors, allowing adaptive radiations (Grant and Grant, 2019; Shimizu-Inatsugi et al., 2017). This mechanism may have played an important role in the diversification of *Sauvagesia* cladeA, shaping their highly specialised morphology to fire-prone and high drought environments, nutrient-deficient soils, and topographically complex landscape of the rocky outcrops along the Espinhaço Range (Giulietti and Pirani, 1987), as suggested by other mountain lineages (*e.g.,* Hughes and Eastwood, 2006; Paule et al., 2017).

### 4.3 Phylogenetic relationships within Sauvagesieae in the light of a complex molecular evolution

Despite the phylogenomic complexity found in Sauvagesieae, we resolved the main clades of the tribe and found evidence of evolutionary processes including incomplete lineage sorting, gene and genome duplication, and reticulation across clades. The uncertainty about the phylogenetic placement of *Tyleria* is worth highlighting. Concordance factors could also help to distinguish between the two probable causes of low resolution in this genus: (i) lack of informative data (*e.g.,* fragmentary sequences), limiting our ability to resolve single locus trees, or (ii) many of the loci contain genuinely conflicting signals. We expect similar values of gCF and sCF when the only phenomenon causing low quartet support is genuine discordant signal among gene trees (Minh et al., 2020). According to the moderate to high sCF values for some couples of species (Fig. S17), there is enough data to inform the relationships within *Tyleria*, but there are many conflicting signals between loci about the position of these couples. This pattern probably comes from processes such as ancient duplication and incomplete lineage sorting. We have evidence for both, the proportion of *Q*2*/Q*3 *≈* 1 for *Tyleria* suggests ILS (Fig. 4), and the intermediate values of AD and LH suggest signals of polyploidisation (Fig. 6a; Table S4), as well as the estimation of ploidy using nQuack, but in most cases with high uncertainty (Table S6). However, *Tyleria* samples showed the lowest median sequence size across genes (Fig. S25), despite the moderate to high number of genes captured and the total sequence length (Fig. 4 and Table S3). Consequently, the weird pattern of the phylogenetic position of this genus could be the result of both a complex history and low-quality sequences (Sayyari et al., 2017). Further efforts are required to better sample this genus and explore duplication and reticulation events, which is challenging because *Tyleria* is a highly restricted genus from the mountain tops of the Amazon-Guyana region such as *Serro da Neblina* tepui (Maguire and Wurdack, 1961; Sastre, 2003).

On the other hand, we corroborated the finding of Schneider et al. (2020) regarding the paraphyly of *Sauvagesia*, which was divided into two distinct clades based on a broader genetic and taxonomic sampling. *Sauvagesia* cladeB includes the type of the genus, *S. erecta* L. as well as representatives of the formerly segregated genera *Pentaspatella*, *Roraimanthus* and *Leitgebia* (Schneider et al., 2014, 2020). This clade is widespread in Central and northern South America except for the pantropical *S. erecta*. *Sauvagesia* cladeA comprises the type of the genus *Lavradia*, *L. vellozii* Vell. ex A.St.-Hil., synonymised under *Sauvagesia vellozii* by Sastre (1971), along with all other species previously described in this genus. Members of this clade are endemic to the *campo rupestre* vegetation in the Espinhaço Range, usually with a highly restricted distribution (Sastre, 1971, 1981; Queiroz-Lima et al., 2023). The distinction between these two clades was initially proposed by Sastre (1971) in a detailed study using numerical taxonomy (Sokal, 1963), where species of the former genus *Lavradia* were recovered as a well-defined group, morphologically characterised by the presence of an internal whorl of staminodes fused in a corona that envelopes the reproductive organs, the absence of calcium oxalate crystals in the leaves, and the lack of an external whorl of free staminodes. Although Sastre (1971) transferred all *Lavradia* names to *Sauvagesia*, as well as the SE Asian *Neckia serrata* Korth., he later recognised the morphological cohesion of the *Lavradia* species and grouped them under the name *Sauvagesia* sect. *Sauvagesia* subsect. *Vellozianae* (Sastre, 1981).

In contrast, the position of *Sauvagesia africana*, sequenced here for the first time, as sister to *Adenarake* + *Sauvagesia* cladeB (but weakly supported, Figs. 4, S7-S17) contradicts the hypothesis proposed by Sastre (1971) about the closer relationship between *S. africana* and all *Lavradia* species, based on the terminal inflorescence and the absence of external staminodes. This interpretation was further supported by the morphology-based cladistic analysis of Amaral (1991). However, *S. africana*, originally described as *Vausagesia africana* Baill., has internal staminodes connate just at the base forming a pseudocorona, distinct from the corona-like structure formed by the entirely fused internal staminodes, characteristic of *Lavradia* (*Sauvagesia* cladeA). A comprehensive taxonomic revision, including estimation of trait evolution, is currently ongoing to elevate the existing subsections of *Sauvagesia* to the genus level, and to morphologically circumscribe the genera *Lavradia*, *Vausagesia*, *Sauvagesia*, and *Adenarake* based on synapomorphies that reflect the phylogenetic history evidenced here.

By increasing molecular data and taxonomic sampling, we also corroborated and improved the support for the relationship between *Blastemanthus* and the clade comprising the *Cespedesia* clade and the African genus *Fleurydora*, as previously suggested by Reinales and Parra-O. (2020) and Schneider et al. (2020). However, the phylogenetic position of *Krukoviella* remains poorly supported with a high gene-tree discordance, probably due to hybridisation as suggested by both HybPhaser (Table S4) and SNaQ (Fig. 6b), instead of limited information given the similar values of the concordance factors (Fig. S17), and the good recovery for this species (Fig. 4 and Table S3). Hybridisation appears to be a widespread process causing phylogenetic conflict and low support within the *Cespedesia* clade. This is also the case within *Rhytidanthera*, where intermediate morphologies between *R. splendida* and *R. sulcata* in the northern Eastern Cordillera of Colombia, and between *R. splendida* and *R. regalis* in the eastern Andean-Amazonian foothills, along with high frequencies of fruits with aborted ovules, support this hypothesis (Reinales *pers. obs.*). Similar patterns have been observed in other Andean plant genera such as *Lupinus* (Nevado et al., 2018), *Oritrophium* (Quinlan et al., 2025), *Senecio* (Dušková et al., 2017; Salomón et al., 2025), among others, prompted by changes in habitat connectivity during Pleistocene glacial cycles. Given the limited number of taxa supported by SNaQ, our results remain inconclusive with respect to specific reticulation events. A focused phylogenetic network analysis for this clade is therefore required to also date these events and improve our understanding of the evolutionary dynamics in this clade.

Within the SE Asian clade, we improved resolution and support for the poorly resolved relationships among *Indosinia*, *Indovethia*, *Neckia*, and *Euthemis* reported by Schneider et al. (2020, Figs. 2 and S2), regardless of the filtering strategy (Fig. S7). This clade showed about 30% of the gene and site concordance factors, and relatively low conflict among gene trees (Q1=76). However, the phylogenetic position of *Neckia* within the clade remains weakly supported. Due to the low difference observed between the gCF and sCF values, and the topological discordance between ML and ASTRAL species tree recovered in this study, as well as by Schneider et al. (2020, 2021), the uncertainty of the relationships of *Neckia* may reflect genuine discordance between gene trees rather than insufficient data (Minh et al., 2020). No evidence of reticulation was found (Fig. 6b-c), but ploidal-level inference for some samples within this clade showed high uncertainty between diploid and tetraploid (Table S6), but with low allele divergence and locus heterogeneity (Table S4). This pattern is consistent with an ancient rather than a recent duplication event, followed by ILS within the clade. Although distinguishing ancient polyploidy, via allo- or autopolyploidy, from ILS is particularly challenging (Meleshko et al., 2021), even more so when using target-capture data, gene-tree discordance caused by ILS is common when internodes are short due to rapid lineage diversification (Koenen et al., 2020; Cai et al., 2021; Dong et al., 2022). Such conditions were found in the deeper nodes of the SE Asian clade (Fig. S17; Schneider et al., 2020, Fig. 2; Schneider et al., 2021, Fig. 3).

Across the species tree, certain samples of the same species such as *Sauvagesia amoena* within *Sauvagesia* cladeB, *S. spicata* and *S. ribeiroi* within cladeA, and *Rhytidanthera sulcata* within the *Cespedesia* clade, were not recovered as clades (Fig. 4). This phylogenetic incongruence is probably due to genuine discordance among gene trees, caused by biological processes such as reticulation followed by a random sorting, sequencing or assembling of different copies (true paralogs), as previously discussed. In contrast, cases such as *Schuurmansia henningsii* Hoogland8954 and *Neckia serrata* Duangjai47 within the SE Asian clade, that pattern is more probably explained by methodological artefacts. Although dual hybridisation allows integration of data from multiple sources (Hendriks et al., 2021, 2023), the little or complete absence of overlap in captured loci across samples can lead to conflicting quartet signals, reducing support values for those branches (Mirarab et al., 2014). Consequently, samples from a single hybridisation, such as those mentioned, may behave as rogue taxa in the species tree. Furthermore, when using summary methods such as weighted ASTRAL, loci with low bootstrap support (*e.g.*, A353; Figs. 3 and S18) contribute less to the final species tree (Fig. S19; Zhang and Mirarab, 2022b). However, unweighted methods are more susceptible to GTEE and generally show lower accuracy, particularly under conditions of high gene tree discordance and moderate to high ILS (Zhang and Mirarab, 2022b). Despite lower support and higher discordance among gene trees derived from the universal probe set compared to the Ochnaceae-specific set, their integration improved both locus and taxon sampling, and the resolution at certain nodes of the species tree (Figs. S19-21). These findings highlight that different gene sets have different resolution power depending on the evolutionary dynamics of each lineage (Chen et al., 2015), and the way each probe set was designed (Siniscalchi et al., 2021; Johnson et al., 2019).

## 5 Conclusion

Target capture sequencing is a cost-efficient way to obtain large amounts of nuclear data for phylogenomic analyses, especially when using a dual hybridisation approach and a predominantly herbarium-based sampling. However, one of the main challenges of target capture data lies in the identification of orthologs, and the difficulty of independently teasing specific sources of gene tree discordance accounting for these factors in phylogenetic inference, even more so in the presence of highly fragmentary material and molecular complex groups. The effect of removing entire genes and samples for paralog warnings in reducing the gene tree estimation error depends on the quality of the dataset and the complexity of the molecular evolution of the group under study. In certain empirical datasets, including the one analysed in this study, this approach did not result in a significant positive effect. However, a comprehensive examination of the data employing multiple filtering strategies, different approaches for paralog assessment, and additional methods such as phylogenetic networks, and ploidy estimation enabled the inference of a robust phylogenetic hypothesis for Sauvagesieae, improving the resolution and support within the SE Asian clade and *Cespedesia* clade, as well as increasing the evidence supporting the paraphyly of *Sauvagesia*.

Beyond improving phylogenetic relationships within Sauvagesieae, identifying consistently recovered clades across different datasets, and highlighting focal clades that required further investigation, this study also contributes to the understanding of molecular evolution in the tribe by exploring probable causes of phylogenetic tree discordance, such as genome duplication and hybridisation. Although the analysis of allele divergence and frequencies, and locus heterozygosity derived from target-capture data provides a novel approach to infer processes such as recent polyploidisation without requiring fresh material and chromosome counts, these metrics can vary depending, for example, on the data, sequencing quality, and the study group. Combining approaches such as chromosome counts and flow cytometry to test the accuracy of nQuack models in Ochnaceae is desirable, as well as producing high-quality genome assemblies to test ancient genome duplication events, as they have a higher impact on phylogenetic discordance and lack of resolution. More detailed and clade-specific network analyses on phasing haplotypes are also desirable for *Sauvagesia* cladeA, *Sauvagesia* cladeB, and *Cespedesia* clade, in order to further investigate the role of hybridisation in the diversification of these mountain lineages.

## Authorship contribution statement

S.R., J.R.P. and F.F. conceptualised the study, with contributions by G.A.B. and A.R.Z. S.R., J.R.P. and F.F. provided financial support. S.R. carried out herbarium, field, and wet laboratory work with support from A.R.Z. and F.F. S.R. conducted formal analyses with contributions by G.A.B. in phylogenetic network inference. S.R., Do.Cardoso and Da.Cardenas contributed with samples, advised on taxonomy and morphology, and contributed taxonomic expertise on Ochnaceae. S.R. wrote the first draft and produced the figures. All co-authors improved the manuscript, contributed to the discussion, and read and approved the final version of the manuscript.

## Acknowledgements

We would like to thank curators and supporting staff for herbaria visited during this project: Jacek Wajer (BM), Carlos Parra (COL), Luciano Paganucci (HUEFS), Sue Zmartzy (K), João Stehmann (MG), Renato Mello-Silva (SPF), and Germinal Rouhan (P), and the following colleagues for their support in obtaining samples: Annelise Frazão, Danilo Zavatin, Matheus Colli-Silva, Luana Sauthier, Roberto Baptista, Renato Ramos and Maria Costa. We are grateful to Jaqueline Alves, Aline Possamai, João Monzoli, Daniela Zappi, Nigel Taylor, Douglas Henrique, Thais Vasconcelos, Raquel Pizzardo and Miriam Antonicelli for their support during the fieldwork. We acknowledge the valuable assistance in the molecular wet laboratory of Robyn Cowan, Catherine McGinnie, Laszlo Csiba, and Paulo Gaem from RBGK, and Matheus Colli-Silva and Adriana Marchioni from SPF, as well as Carlos Parra, José Murillo, and Janice Valencia for valuable insights to yield viable DNA extracts for difficult samples. Many thanks also to Juan Viruel, Sara Martin, George Tiley and Cássia Bitencourt for useful insights and discussion about bioinformatics, and Julio Schneider and Toral Shah for valuable information about previous analyses in Ochnaceae. The development of this project was also made possible by the StackOverflow.com community and the many tutorials and scripts shared on GitHub by Sidonie Bellot, Liam Revell, and Matt Johnson. We also thank Carolina Siniscalchi and Marcelo Reginato for their useful comments on the first version of this manuscript.

## Data Availability

Raw sequence data produced for this study have been deposited in the European Nucleotide Archive (ENA) at the European Bioinformatics Institute (EMBL-EBI) under accession number PRJEB101313 (https://www.ebi.ac.uk/ena/browser/view/PRJEB101313). Supplementary figures and tables are available in the article as online supplementary material. Data and code underlying this study are available in Zenodo, at https://doi.org/10.5281/zenodo.17538083). Custom scripts for formal analyses are also available at https://github.com/spreinalesl/sauvagesieae_phylogeny.

