## supplementary material for "Integrating lineage-specific and universal genomic probes illuminates phylogenetic relationships and molecular evolution in Sauvagesieae (Ochnaceae)"

### 1 Methods

#### 1.1 DNA extractions and library preparation

Different protocols for DNA isolation have been tested including DNeasy Plant Mini Kit (QIAGEN) following manufacture procedure, DNA isolation with Sodium Dodecyl Sulphate-SDS (Mogg and Bond, 2003), standard Cetyltrimethylammonium bromide (CTAB) protocol (Saghai-Maroof et al., 1984; Doyle and Doyle, 1987; Doyle and Dickson, 1987), Sorbitol-based wash buffer and high-salt CTAB (Tel-Zur et al., 1999; Souza et al., 2012; Inglis et al., 2016). In addition, multiple variations of these protocols were also tested including different amounts of $\beta$mercaptoethanol, Polyvinylpyrrolidone - PVP40 and Proteinase K, along with variations in leaf storage (*i.e.*, silica-dried and herbarium), amount of initial plant tissue, and types of tissue disruption. Regardless of the protocol used, the retrieved DNA concentration did not exceed $10ng/\mu$ in mean for most samples of *Sauvagesia* and *Tyleria*, and samples purity assessed via the ratios between absorbance values of 260nm vs 280nm ($A260/A280$) and the 260nm vs 230nm ($A260/A230$), were also out of the desired values (Figure S1). Leaves of those genera probably contain high quantities of polyphenols, tannins, and polysaccharides hindering DNA isolation of good quality. In contrast, DNA extractions from *Ochna serrulata* samples produced good concentrations and quality using all protocols. Finally, DNA was extracted using a modified CTAB protocol (Saghai-Maroof et al., 1984; Doyle and Doyle, 1987; Doyle and Dickson, 1987), including a wash step with CTAB 10% + NaCl buffer between two cycles of extraction with chloroform:isoamyl alcohol $24 : 1v/v$. The DNA was precipitated in pre-cooled isopropanol at $-20°C$ for $8 - 10$ days, to maximise the final DNA concentration. The resulting pellet was washed twice with 70% ethanol and one with 96% ethanol, and after room temperature dried, was dissolved in the TE buffer.

#### 1.2 Custom target files

Custom target files resulted in higher capture success and longer sequences compared to the standard target file used by PAFTOL (https://github.com/mossmatters/Angiosperms353/blob/master/Angiosperms353_targetSequences.fasta). We compared the percentage on target, number of genes with sequences, and number of loci with paralogous sequences after sequence assembly using different target files in Hybpiper. In general, the mega353 target file filtered by Malpighiaceae retrieved more sequences per locus and fewer genes containing paralogous sequences (Figure S2). A custom target file to reads assemble from the Ochnaceae-specific probes was designed based on the scaffold sequences used to design the probe set (available in https://zenodo.org/record/4278991). Detailed code to generate target files, and the corresponding files are available in https://github.com/spreinalesl/sauvagesieae_phylogeny.

In order to detect shared loci between both probe sets, we performed a comparison among the 353 exon sequences including in the Angiosperms353 set for *Hypericum perforatum* (BNDE), *Ochna serrulata* (CKDK), *Garcinia oblongifolia* (FWCQ) and *Salix viminalis* (KKDQ), against the scaffold sequences of *Ochna serrulata* (CKDK) using to design the Ochnaceae set of probes. Blast hits with E-value less than 0.01 were considered good hit for homology matches. Among the ten couples of loci tagged as duplicates, we removed one of them from the target file after checking the overlap between Angiosperms353 and Ochnaceae-specific probes. Each of the ten couples was aligned using MAFFT v.7.520 (Nakamura et al., 2018) and observed using UGENE. For 9 of those couples, the a353 sequences are contained in the scaffold sequences, then, the scaffold sequences were maintained in the Ochnaceae-specific target file and removed from the a353 target file. For the couple g2094005-g5943, the a353 sequence has at least 824 bp without overlap, so, the sequences for that gene were maintained in the a353 target file and the correspondent scaffold was removed. Detailed code is available in https://github.com/spreinalesl/sauvagesieae_phylogeny.

### 1.3 Filtering strategies

**Missing data:** After testing different strategies to remove known positions (N) and short sequences, we decided to remove sequences with more than 50% of unknown positions (N), just if the sequence has no known sequences longer than 150 bp between unknown positions (Figure S3). Those sequences have at least 150 known positions to be aligned, even if more than 50% of their total sequence correspond to unknown positions. After removing those sequences for each locus, sequences with less than 25% of the median sequence length for that locus, or shorter than 150 bp, whichever was longer, were also removed.

**Paralogs assessment:** For the dataset *inform_paralogs* (dataset v) all genes and samples including all paralogs for each gene were retrieved using the Hybpiper script `paralog_retriever.py`. Sequences were aligned using the `auto` option of MAFFT which automatically selects an appropriate strategy according to data size. Trees were inferred using an approximatelymaximumlikelihood approach implemented in FastTree2 (Price et al., 2010) with GTR as nucleotide substitution model. An automated inspection of the phylogenetic grouping of all paralogs for each sample was implemented in a custom script using the function `is.monophyletic` of the R package `ape v.5.7.1`. We kept the *.main* sequence for samples for which all competing contigs (paralogs) formed a clade, as there is no evidence of paralogy (Figure S4, arrowhead). When competing contigs for a sample were reconstructed in different clades, the entire sample was removed from that locus (Figure S4, asterisks).

### 2 Figures

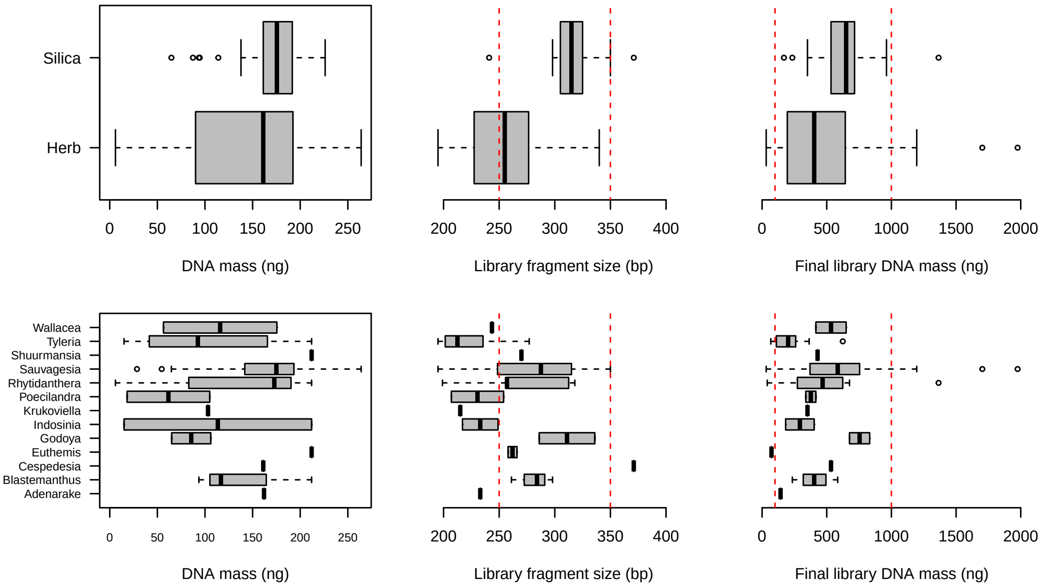


Figure S1: Summary of DNA extractions and library preparation for 96 Sauvagesieae samples. Red lines represent acceptable values.

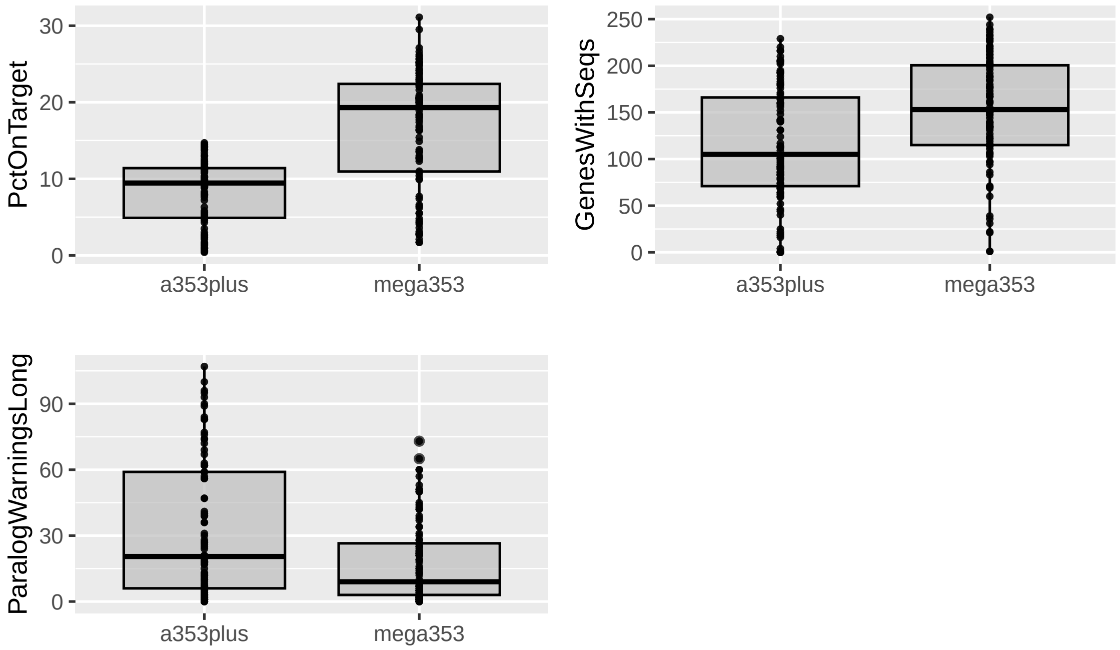


Figure S2: Comparison of the sequence assembly using Hybpiper on a subset of 80 samples. Two different target files were used for reads mapping and locus assembling, the standard PAFTOL target file (the nucleotide version) - a353plus, and the mega353 target file (McLay et al., 2021) filtering for Malpighiales.

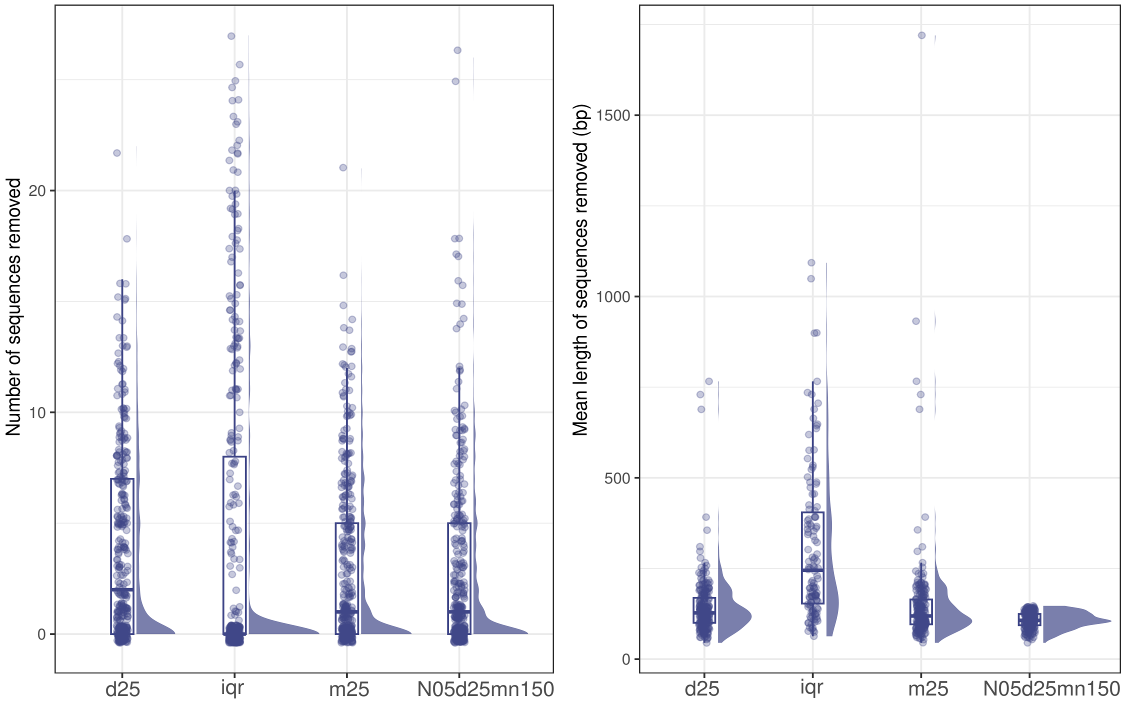


Figure S3: Summary of filtering strategies to remove missing data. Number and mean length of removed sequences after filtering were compared among strategies. We establish the minimum sequence length allowed using the following criteria (from left to right): sequence length <25% median seq length (d25), sequence length shorter than the third quartile plus 3 IQR of the seq length (iqr), sequence length <25% mean seq length, sequence length <25% median seq length or shorter than 150 bp. The last strategy minimises the number of sequences removed, but removes shorter sequences, reducing the chance to remove long and "good" sequences which can be aligned.

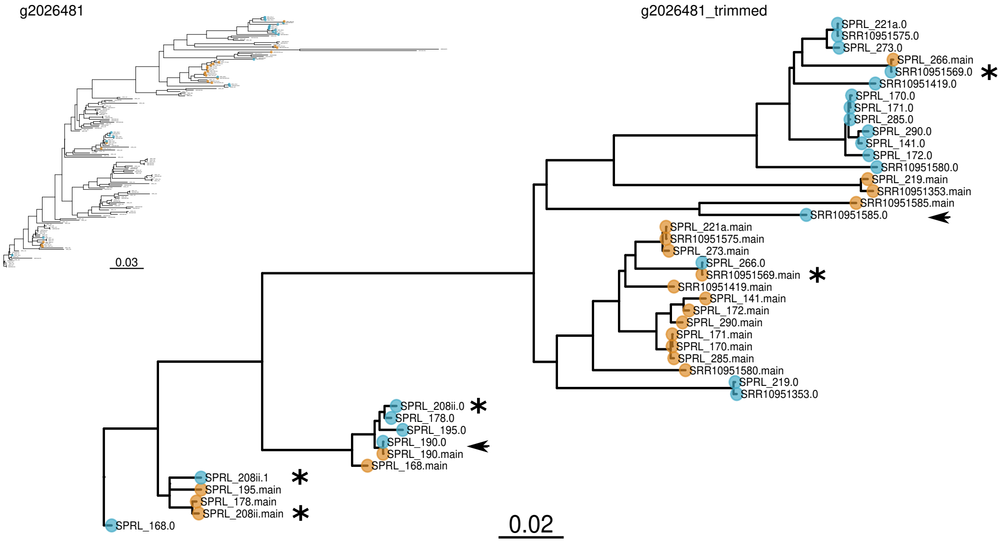


Figure S4: Gene tree for gene g2026481 including all competing contigs per sample retrieved by the Hybpiper script `paralog_retriever.py`. Competing contigs are labeled as *.main* (the largest contig with the greatest percent identity to the reference), and *.0, .1*, etc, for shorter contigs which have at least 85% of the reference sequence and 10X of coverage depth. Arrowheads show contigs for the same sample which form a clade. Asterisks show some examples of competing contigs which were retrieved in different clades, and consequently were removed from this locus.

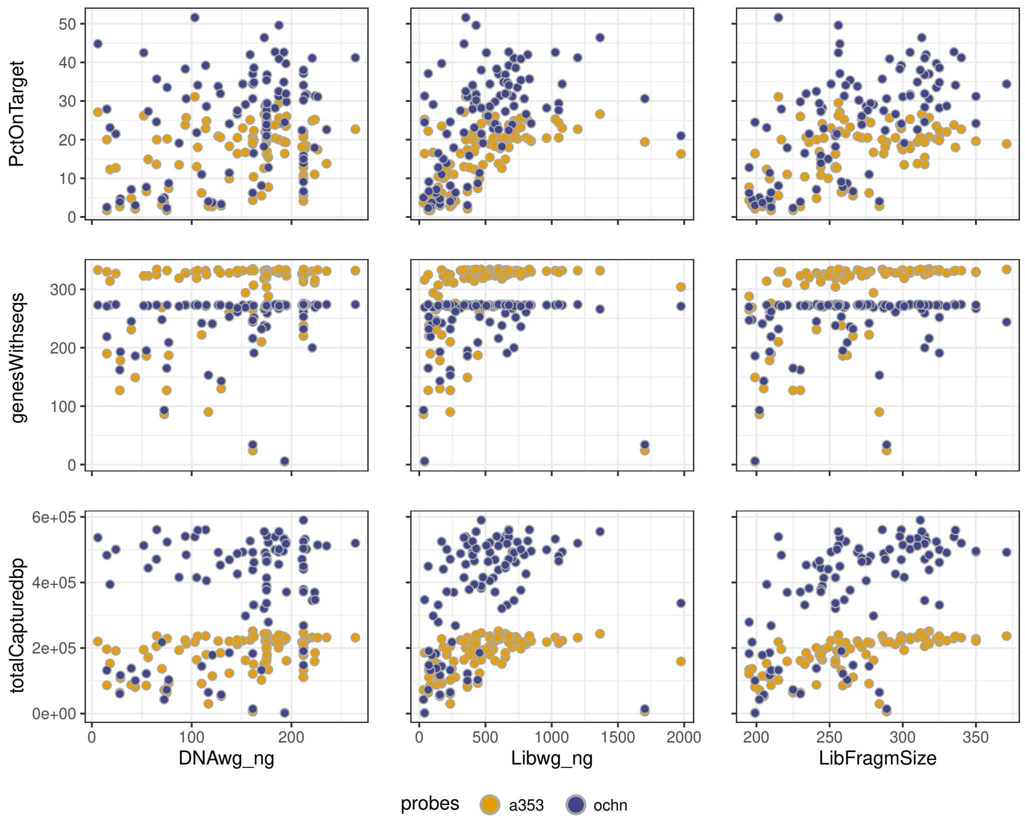


Figure S5: Correlation between library preparation process and different variables used to summarise capture efficiency. Targetcapture efficiency using the universal Angiosperm353 and the Ochnaceae specific probe sets on 96 samples corresponding to 85 species within Sauvagesieae. From left to right: Initial gDNA concentration (ng) after purification, Library concentration (ng), Library fragment size (bp).

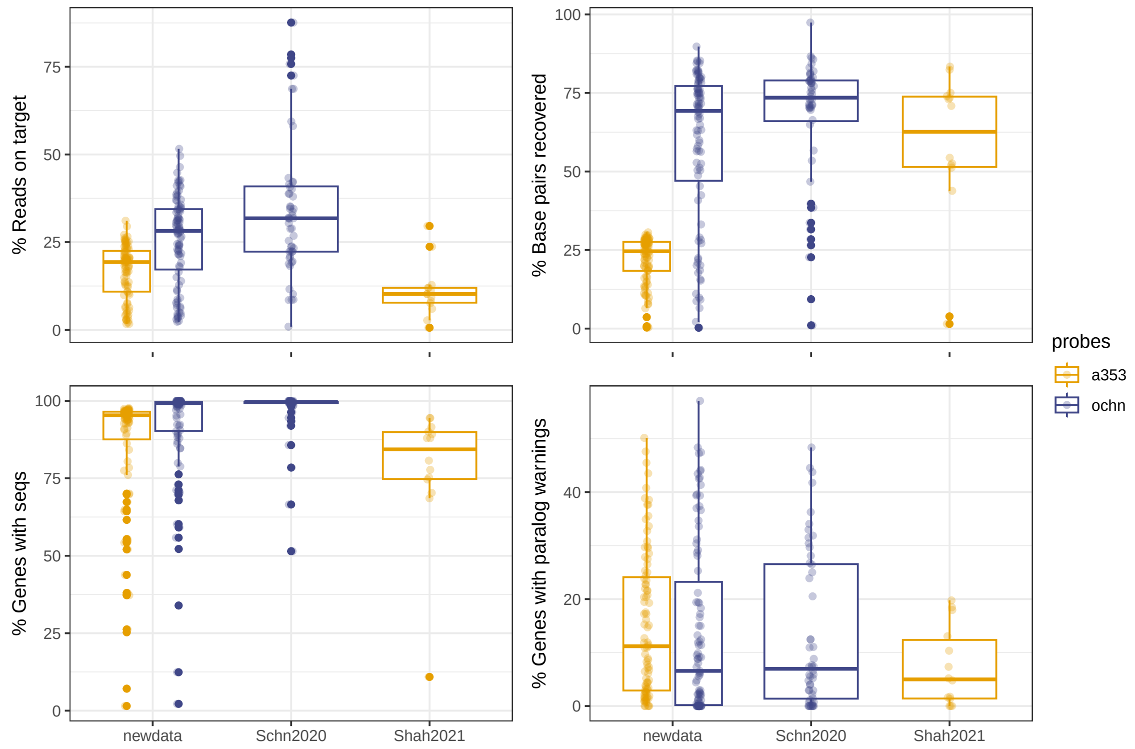


Figure S6: Comparison of the capture efficiency between already published data by Shah et al. (2021) and Schneider et al. (2021) and data produced for this study for Sauvagesieae samples.

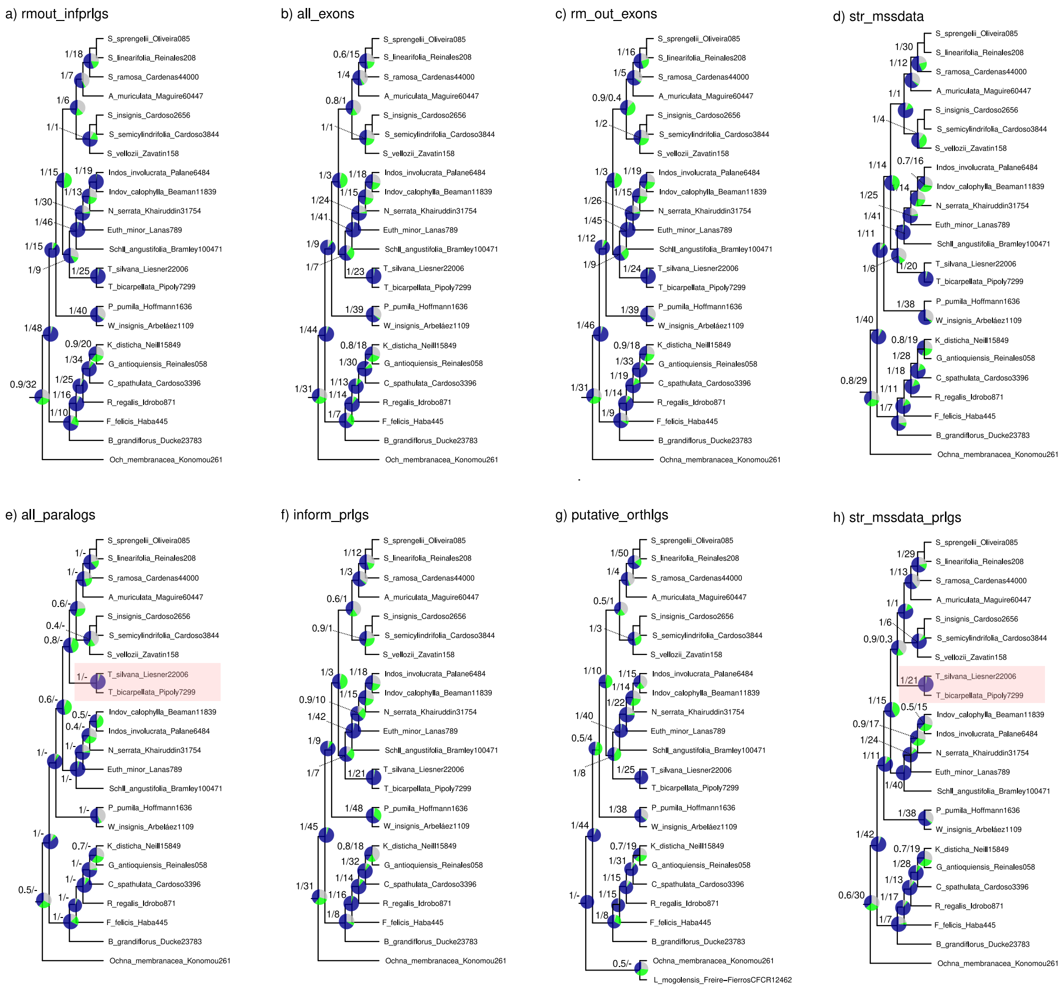


Figure S7: Phylogenetic backbone cropped from the species trees inferred using ASTRAL on the eight datasets resulted from different filtering strategies. Topologies show the main clades within Sauvagesieae. Pie charts indicate conflicting gene trees as quartet support, where blue represents the proportion of input gene trees satisfied by the species tree (Q1). The higher this number, the less discordant the gene trees are. Green and grey represent alternative frequent topologies (Q2 and Q3). Similar proportions of Q2 and Q3 suggest a signal of Incomplete Lineage Sorting (ILS), while different proportions of the two alternative quartets suggest a signal of hybridisation or polyploidisation. Values on the branch correspond to Local Posterior Probability for the main topology (LPP1) and gene concordance factir (gCF) calculated using IQ-TREE2

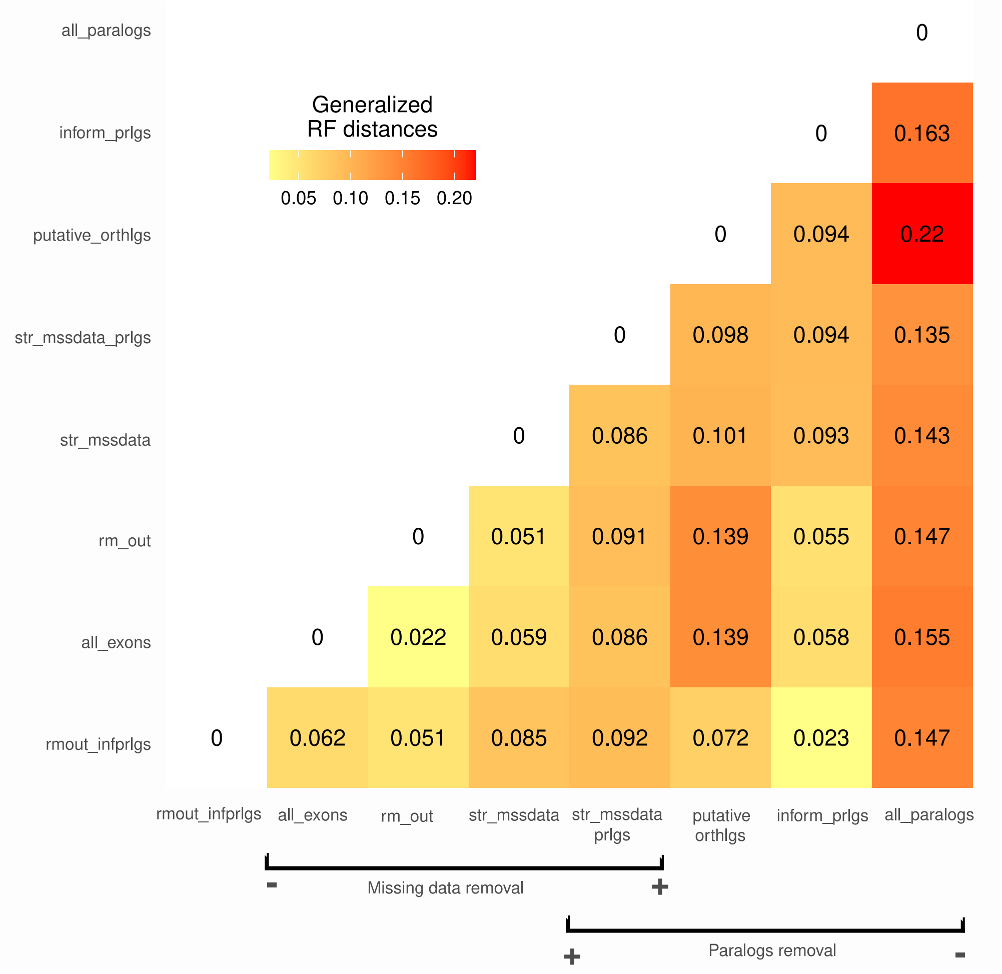


Figure S8: Heatmap of Generalized RobinsonFoulds distances among unrooted species trees estimated with Astral for each dataset. gRF distances were calculated using the function `DifferentPhylogeneticInfo` implemented in the R package `TreeDistance` (Smith et al., 2020). The tree distance is given by the amount of phylogenetic information, *i.e.*, a similarity score, that the pairs of splits share in common, normalized by the sum of the information content of each split in each tree (maximum distance). A distance value of zero represents identical trees, and a distance value of one represents zero similarity; however, zero similarity is uncommon as even the most different trees exhibit some similarity. Tips present in one tree but not in the other are removed before each comparison, because the trees neither hold information in common nor differ regarding these unshared tips.

a) rmout_infprlgs

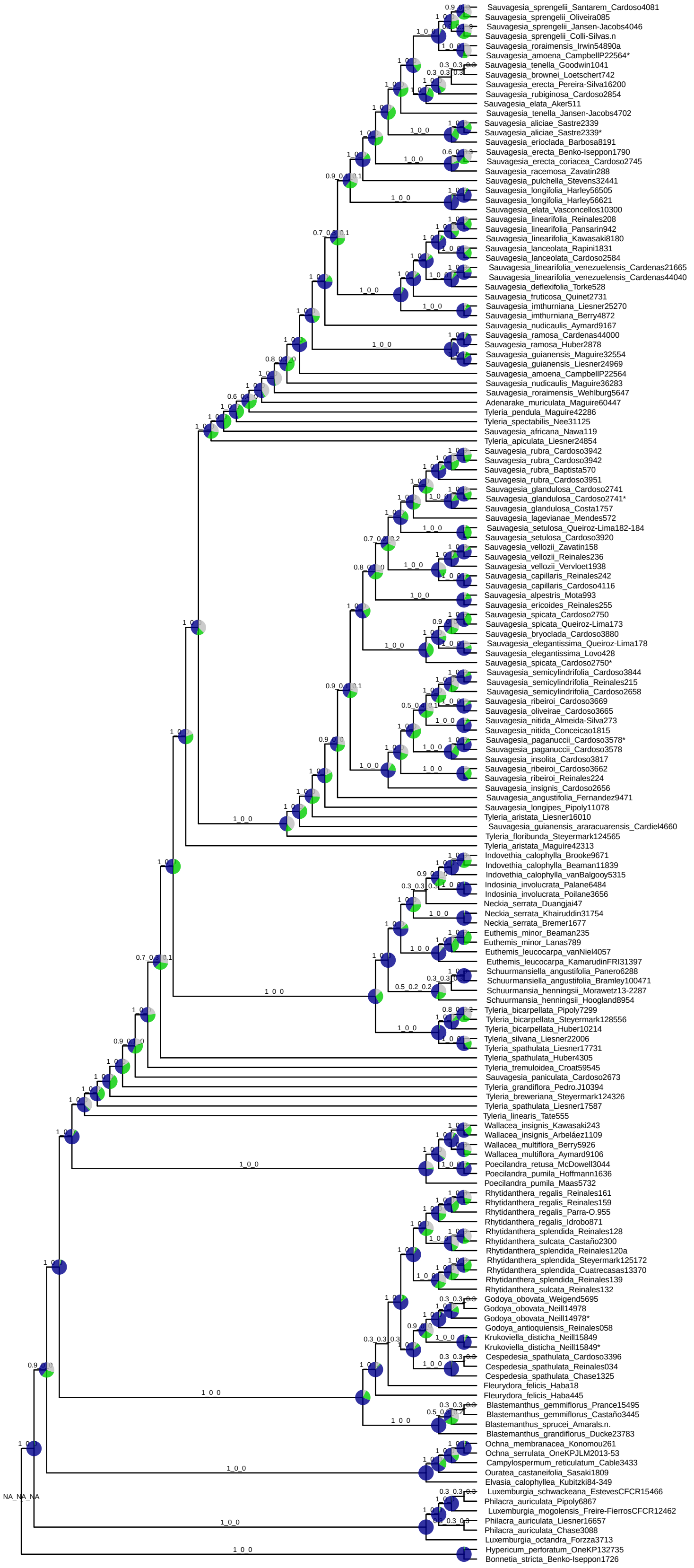


Figure S9: Species tree inferred using Weighted ASTRAL-Hybrid on the dataset rmout_infprlgs. Branch annotations correspond to LPP1_LPP2_LPP3 from ASTRAL. Pie charts indicate conflicting gene trees as quartet support, in blue the proportion of input gene trees satisfied by the species tree (Q1), green and grey represent alternative frequent topologies (Q2 and Q3). Asterisks next to the identical sample pairs highlight the sample sequenced in this study using both probe sets.

b) all_exons

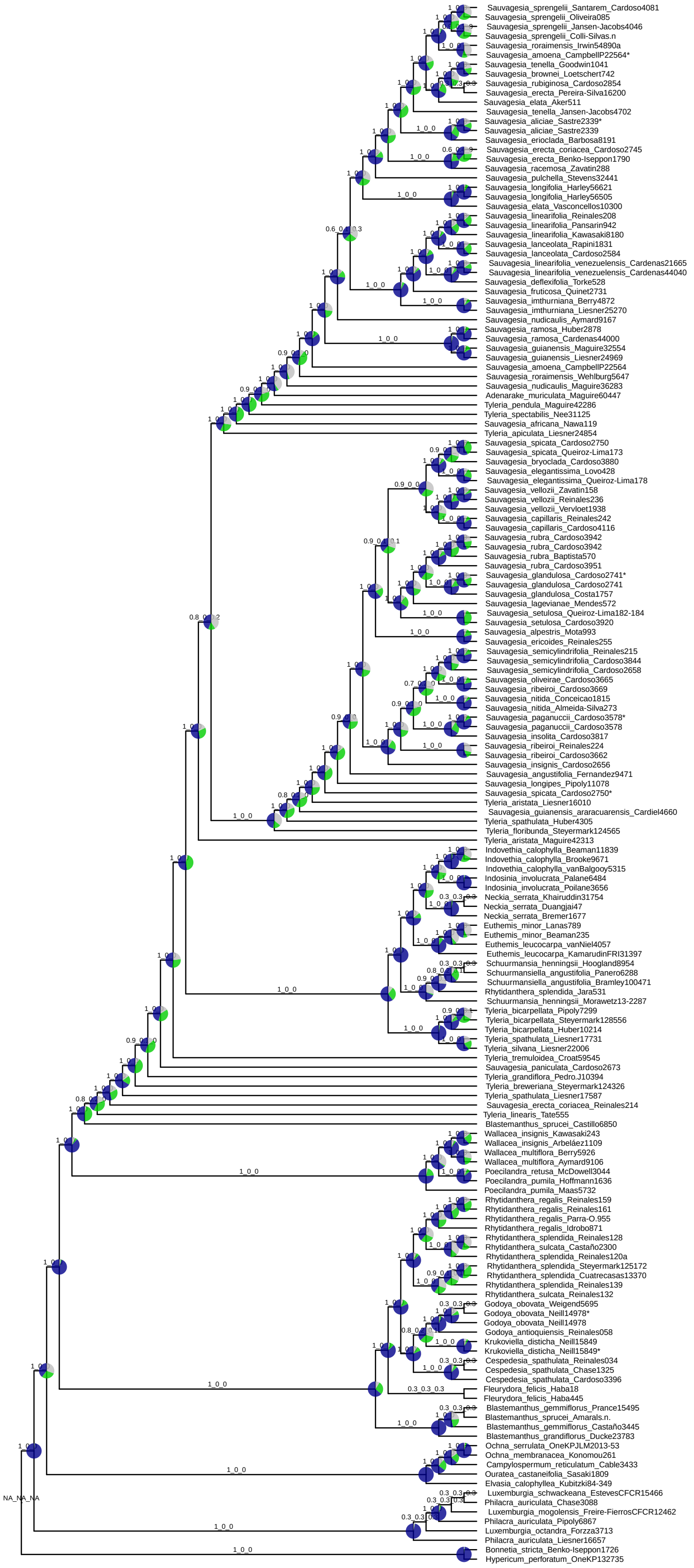

Figure S10: Species tree inferred using Weighted ASTRAL-Hybrid on the dataset all_exons. Branch annotations correspond to LPP1_LPP2_LPP3 from ASTRAL. Pie charts indicate conflicting gene trees as quartet support, in blue the proportion of input gene trees satisfied by the species tree (Q1), green and grey represent alternative frequent topologies (Q2 and Q3). Asterisks next to the identical sample pairs highlight the sample sequenced in this study using both probe sets.

c) rm_out

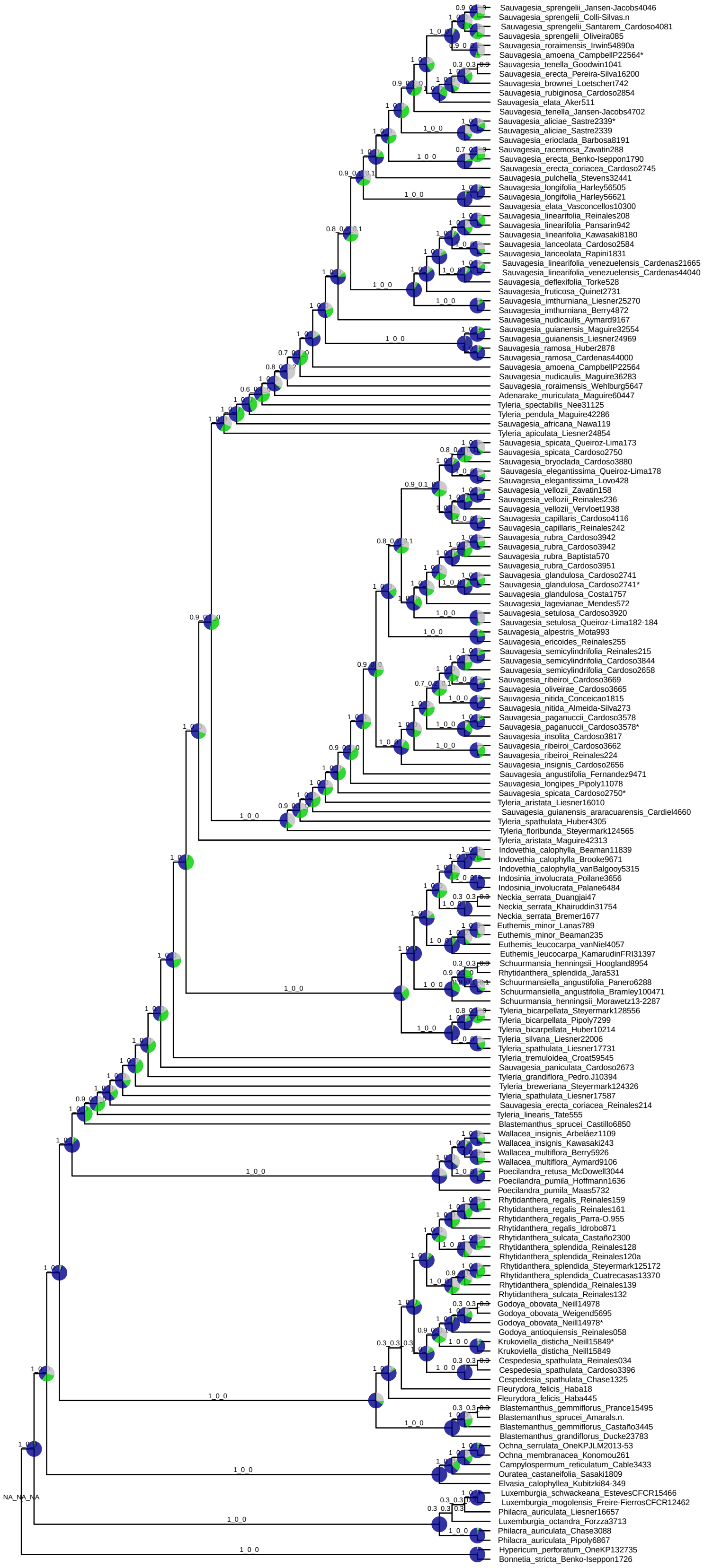

Figure S11: Species tree inferred using Weighted ASTRAL-Hybrid on the dataset rm_out_exons. Branch annotations correspond to LPP1_LPP2_LPP3 from ASTRAL. Pie charts indicate conflicting gene trees as quartet support, in blue the proportion of input gene trees satisfied by the species tree (Q1), green and grey represent alternative frequent topologies (Q2 and Q3). Asterisks next to the identical sample pairs highlight the sample sequenced in this study using both probe sets.

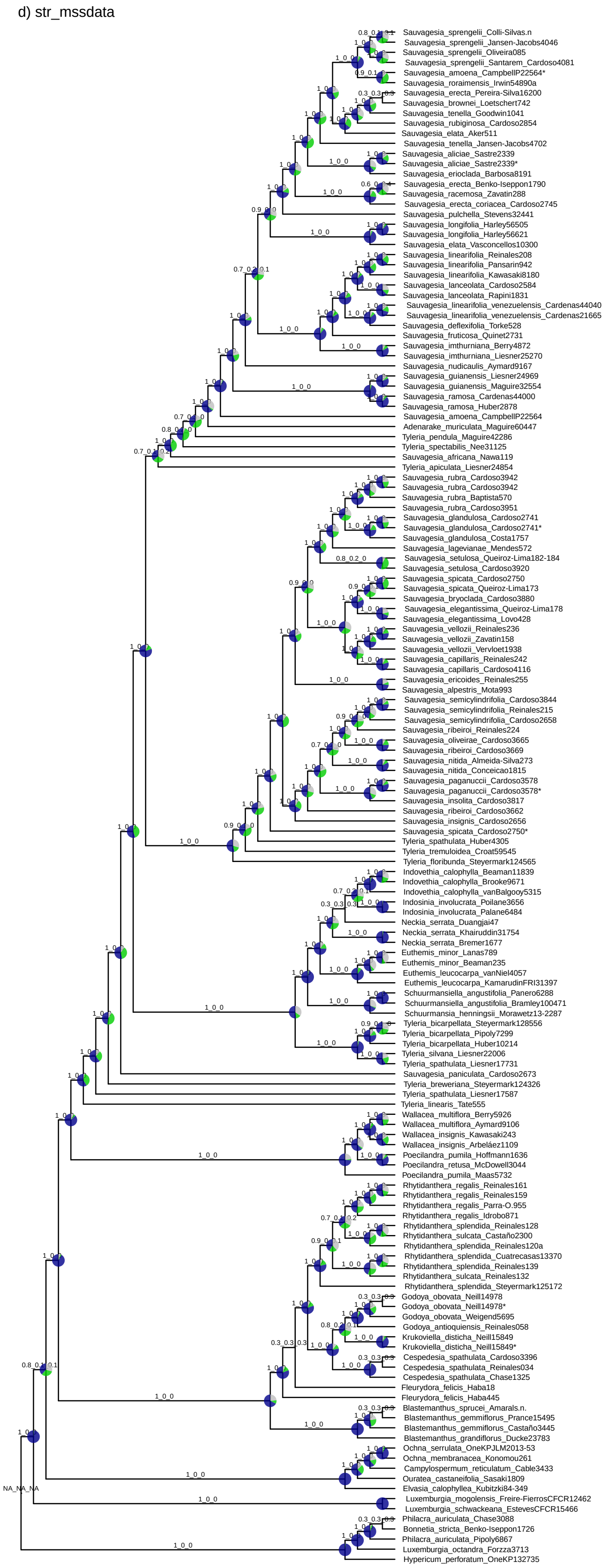


Figure S12: Species tree inferred using Weighted ASTRAL-Hybrid on the dataset str_mssdata. Branch annotations correspond to LPP1_LPP2_LPP3 from ASTRAL. Pie charts indicate conflicting gene trees as quartet support, in blue the proportion of input gene trees satisfied by the species tree (Q1), green and grey represent alternative frequent topologies (Q2 and Q3). Asterisks next to the identical sample pairs highlight the sample sequenced in this study using both probe sets.

e) all_paralogs

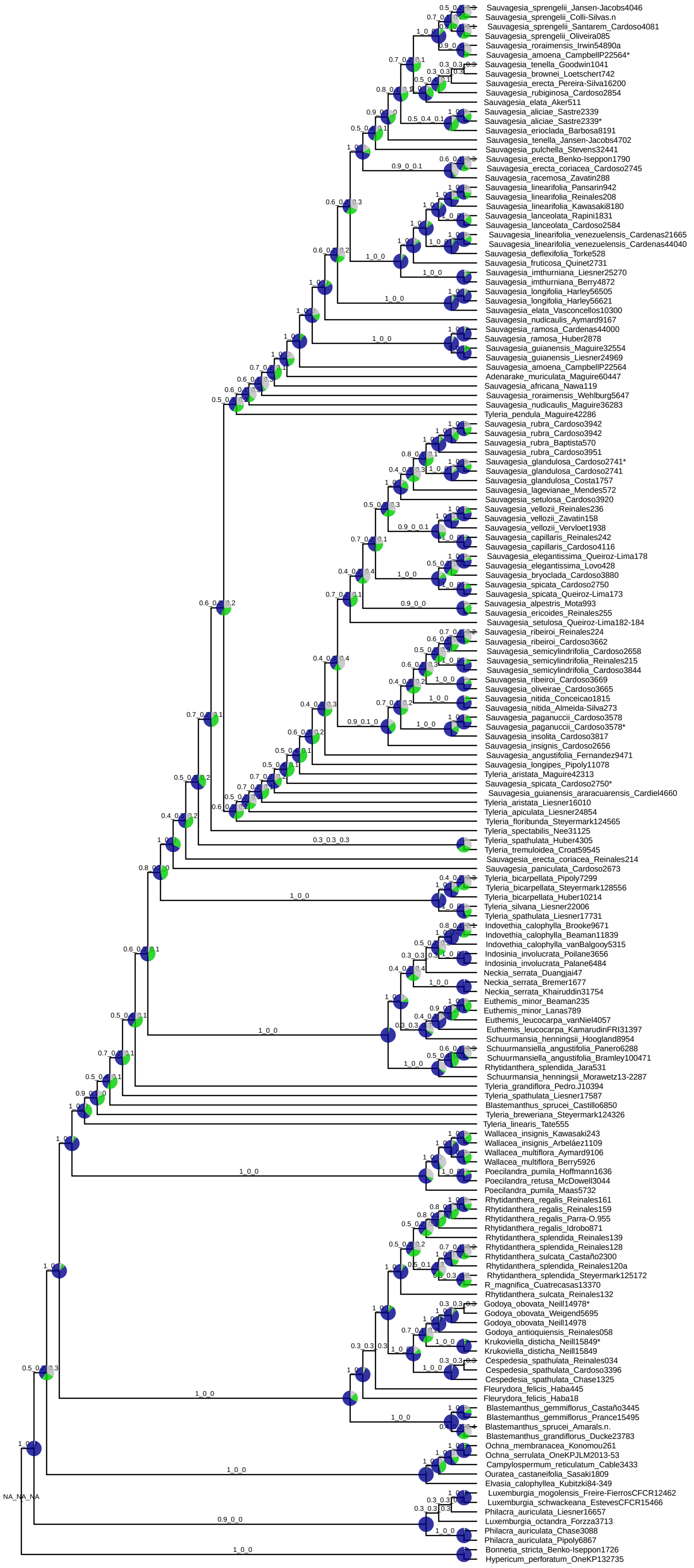

Figure S13: Species tree inferred using ASTRAL-pro2 on the dataset all_paralogs. Branch annotations correspond to LPP1_LPP2_LPP3 from ASTRAL. Pie charts indicate conflicting gene trees as quartet support, in blue the proportion of input gene trees satisfied by the species tree (Q1), green and grey represent alternative frequent topologies (Q2 and Q3). Asterisks next to the identical sample pairs highlight the sample sequenced in this study using both probe sets.

f) inform_paralogs

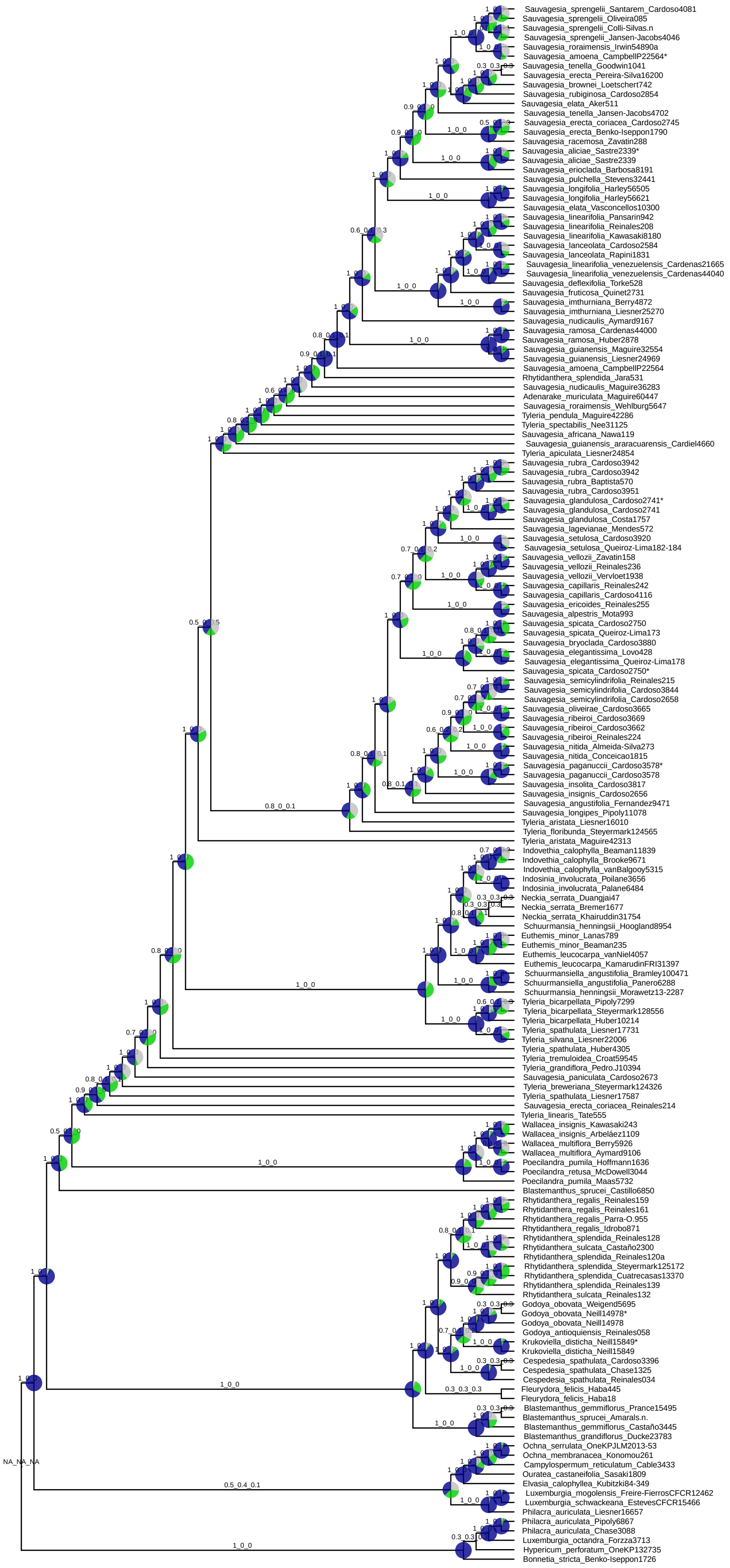


Figure S14: Species tree inferred using Weighted ASTRAL-Hybrid on the dataset inform_prlgs. Branch annotations correspond to LPP1_LPP2_LPP3 from ASTRAL. Pie charts indicate conflicting gene trees as quartet support, in blue the proportion of input gene trees satisfied by the species tree (Q1), green and grey represent alternative frequent topologies (Q2 and Q3). Asterisks next to the identical sample pairs highlight the sample sequenced in this study using both probe sets.

g) putative_orth

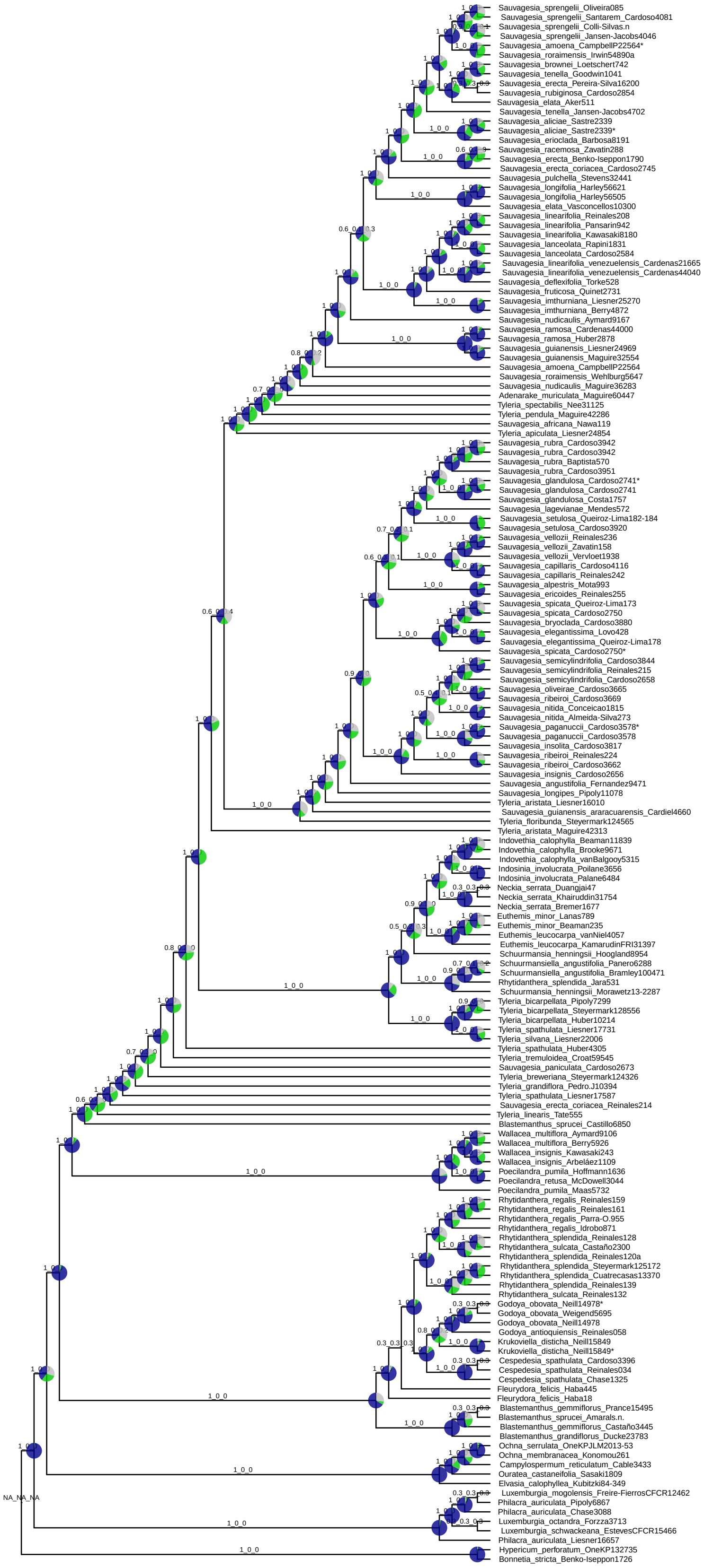


Figure S15: Species tree inferred using Weighted ASTRAL-Hybrid on the dataset putative_orthlgs. Branch annotations correspond to LPP1_LPP2_LPP3 from ASTRAL. Pie charts indicate conflicting gene trees as quartet support, in blue the proportion of input gene trees satisfied by the species tree (Q1), green and grey represent alternative frequent topologies (Q2 and Q3). Asterisks next to the identical sample pairs highlight the sample sequenced in this study using both probe sets.

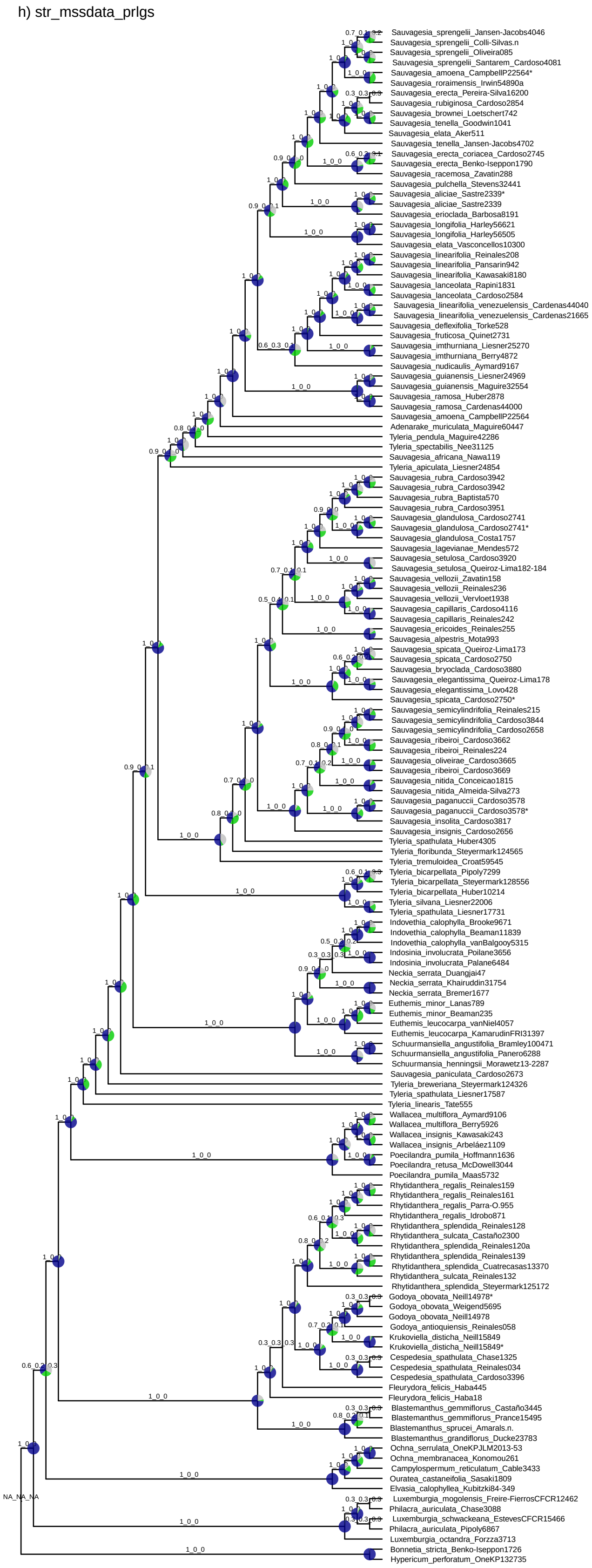


Figure S16: Species tree inferred using Weighted ASTRAL-Hybrid on the dataset str_mssdata_prlgs. Branch annotations correspond to LPP1_LPP2_LPP3 from ASTRAL. Pie charts indicate conflicting gene trees as quartet support, in blue the proportion of input gene trees satisfied by the species tree (Q1), green and grey represent alternative frequent topologies (Q2 and Q3). Asterisks next to the identical sample pairs highlight the sample sequenced in this study using both probe sets.

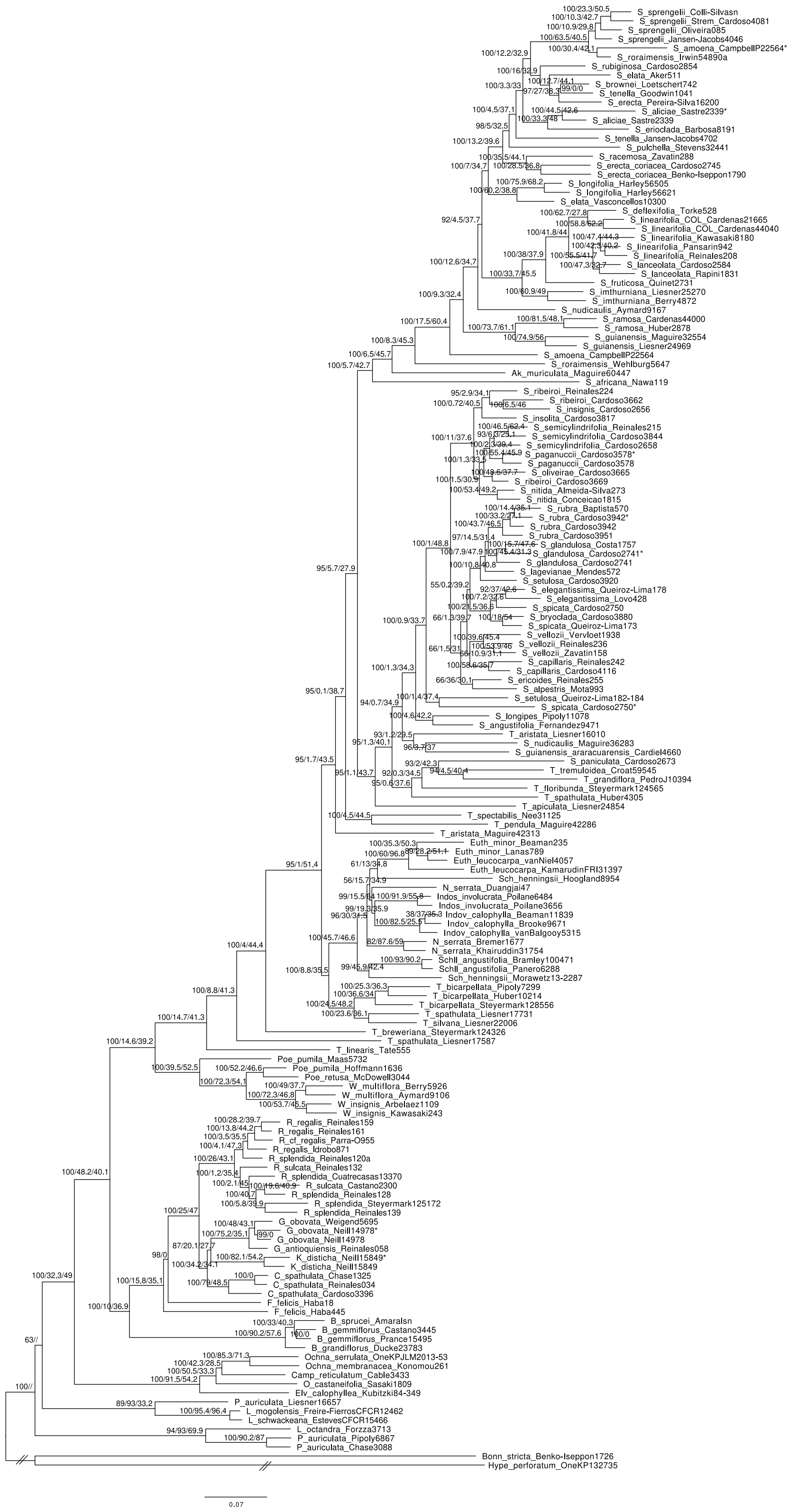

Figure S17: Species tree for the dataset inform_prlgs using IQ-TREE2 based on a concatenate matrix using 608 regions. Branch supports correspond to ultrafast bootstrap, gene concordance factor (gCF) and site concordance factor (sCF). Asterisks next to the identical sample pairs highlight the sample sequenced in this study using both probe sets.

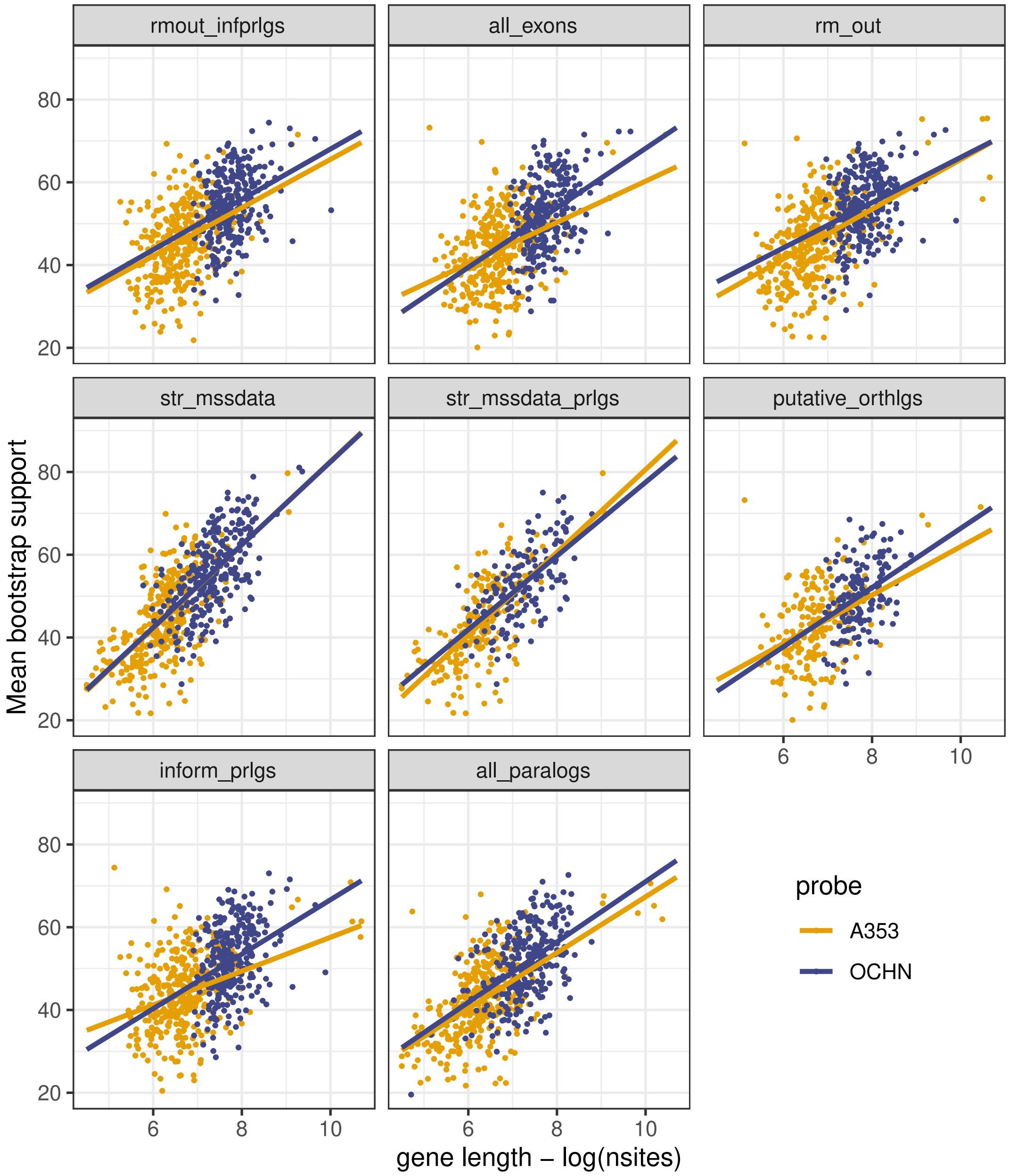


Figure S18: Scatterplot showing the correlation between the gene length and the mean bootstrap support of the gen tree for each probe set. All Filtering strategies showed the same pattern in which the OCHN set has longer genes, and longer genes resulted in gene trees with higher mean bootstrap support.

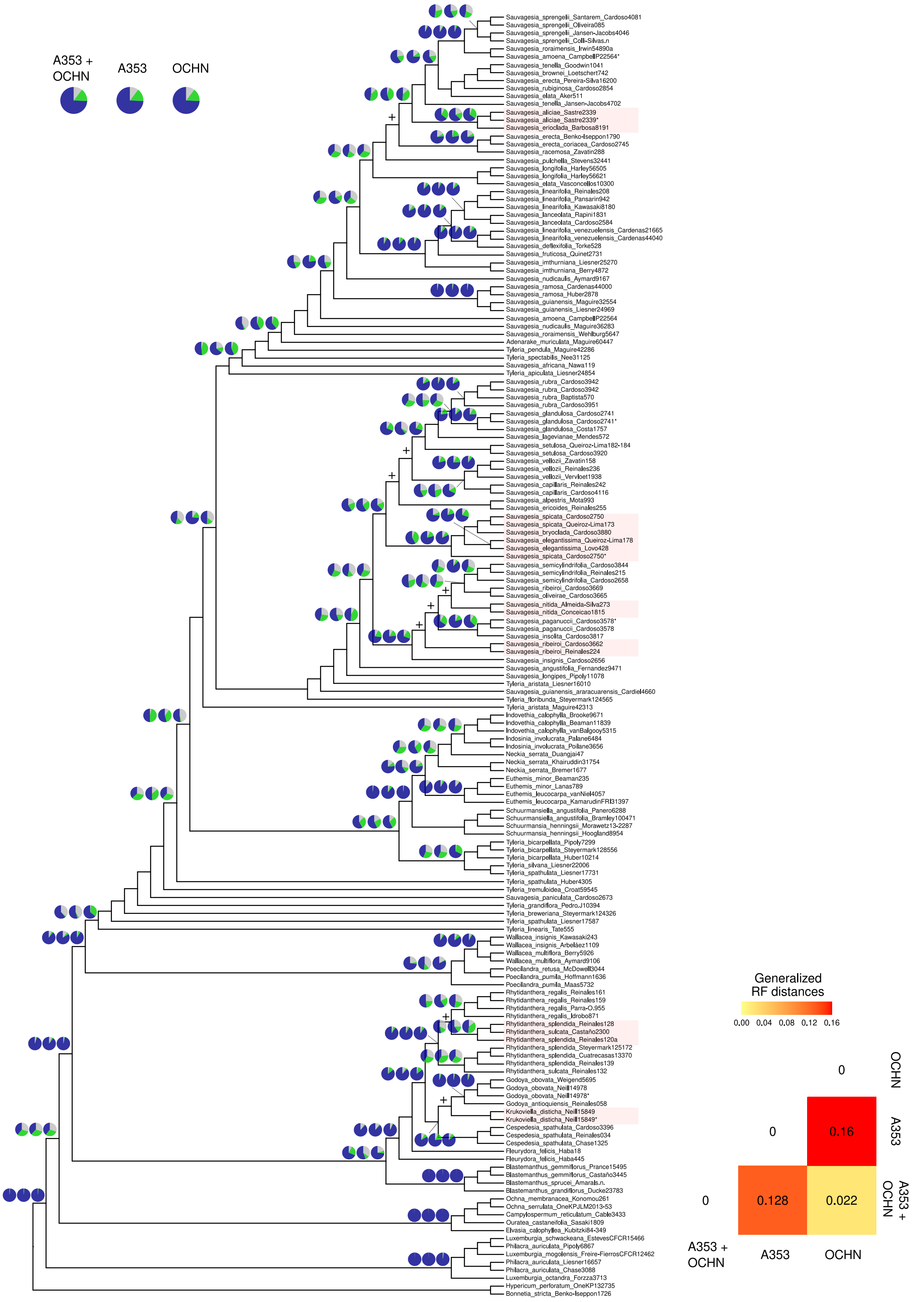


Figure S19: Species tree inferred using Weighted ASTRAL-Hybrid on the dataset rmout_infprlgs. Pie charts indicate conflicting gene trees as quartet support, in blue the proportion of input gene trees satisfied by the species tree (Q1), green and grey represent alternative frequent topologies (Q2 and Q3). The three different pie charts represent independent analysis using both probe sets, and A353 and OCHN genes separately (see Figs. S20 and S21 for respective species trees). Black crosses represent conflicting nodes between the three analyses, and clades with different positions in those analyses are highlighted. Asterisks next to the identical sample pairs highlight the sample sequenced in this study using both probe sets. Topological differences between the three species trees were measured by calculating the generalised Robinson-Foulds distances between trees, as implemented in the R package TreeDist v.2.9.1.

a) A353

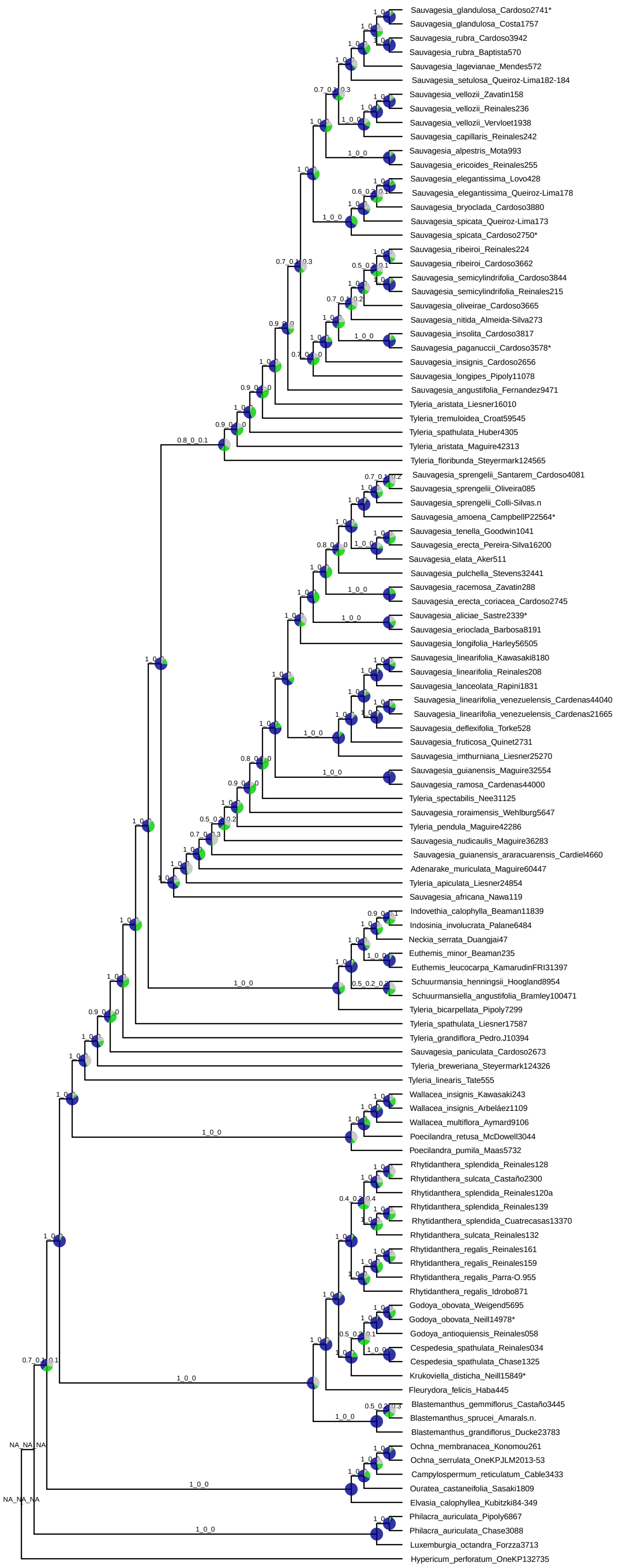


Figure S20: Species tree inferred using Weighted ASTRAL-Hybrid on the dataset rmout_infprlgs using only A353 genes. Branch annotations correspond to LPP1_LPP2_LPP3 from ASTRAL. Pie charts indicate conflicting gene trees as quartet support, in blue the proportion of input gene trees satisfied by the species tree (Q1), green and grey represent alternative frequent topologies (Q2 and Q3). Asterisks next to the identical sample pairs highlight the sample sequenced in this study using both probe sets.

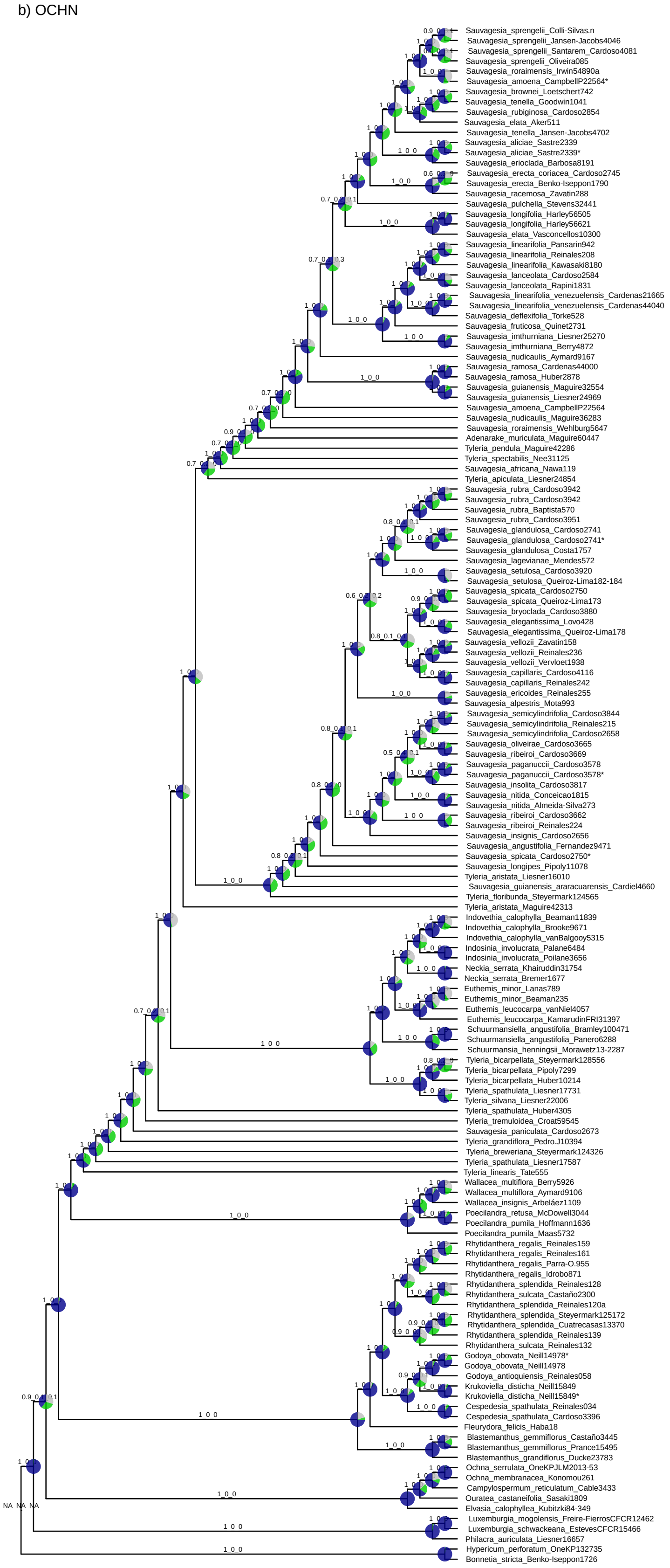


Figure S21: Species tree inferred using Weighted ASTRAL-Hybrid on the dataset rmout_infprlgs using only OCHN genes. Branch annotations correspond to LPP1_LPP2_LPP3 from ASTRAL. Pie charts indicate conflicting gene trees as quartet support, in blue the proportion of input gene trees satisfied by the species tree (Q1), green and grey represent alternative frequent topologies (Q2 and Q3). Asterisks next to the identical sample pairs highlight the sample sequenced in this study using both probe sets.

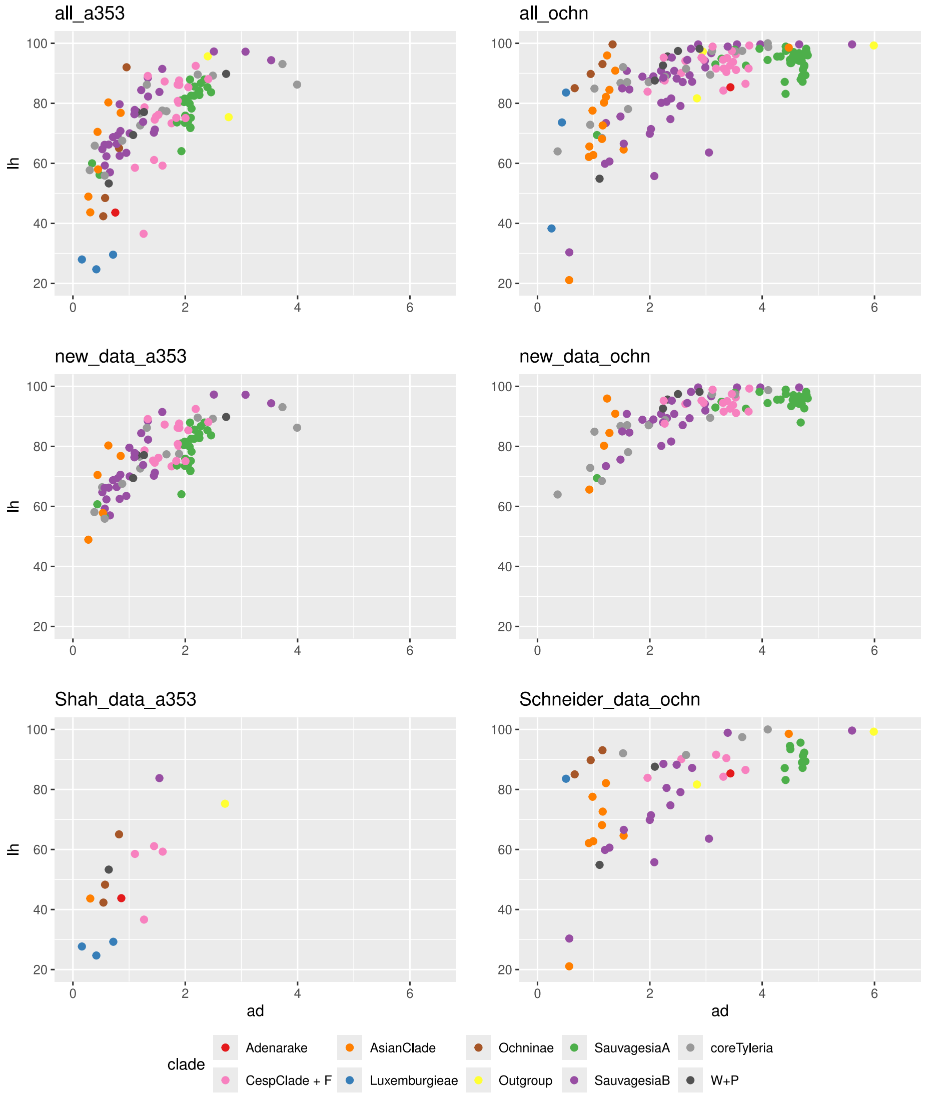


Figure S22: Scatterplot showing the allele divergence (AD) and locus heterozygosity (LH) of all samples after removing highly variable (outliers) samples and genes, and those with less than 10% of sampling. Patterns for previously published and newly sequenced data are compared, as well as patterns displaying for each of the probe sets independently. 'Normal diploids' show roughly $LH < 90\%$ and $AD < 1\%$, while highly and recent polyploids tend to have high AD ($> 3.5\%$) and LH ($> 80\%$) values, and hybrids usually show high levels of LH ($> 90\%$) and intermediate AD values ($1 - 4\%$). AD and LH values are relative and can vary depending on the dataset, sequencing quality, study group, and many other factors.

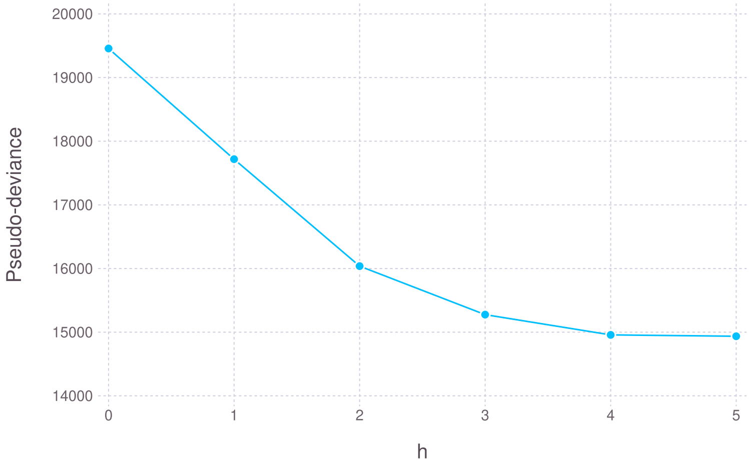


Figure S23: Pseudo-likelihood values for SNaQ independent runs considering the maximum number of hybridisations to be h=0 to h=5.

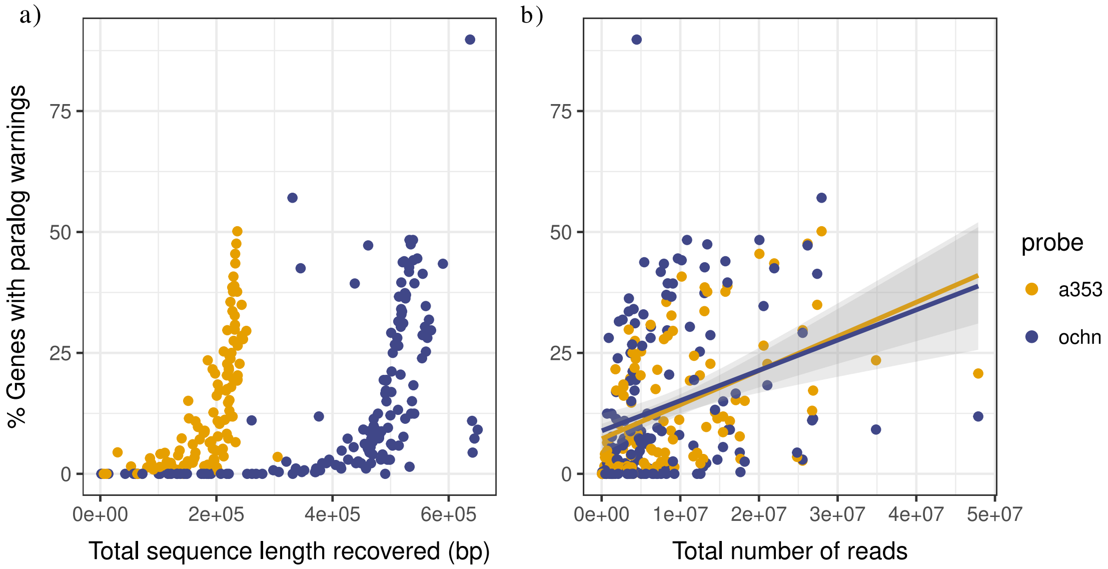


Figure S24: Correlation between capture metrics and the percentage of genes with paralog warnings recovered by Hybpiper. **a)** Total sequence lenght captured per sample. **b)** Number of sequenced reads per sample.

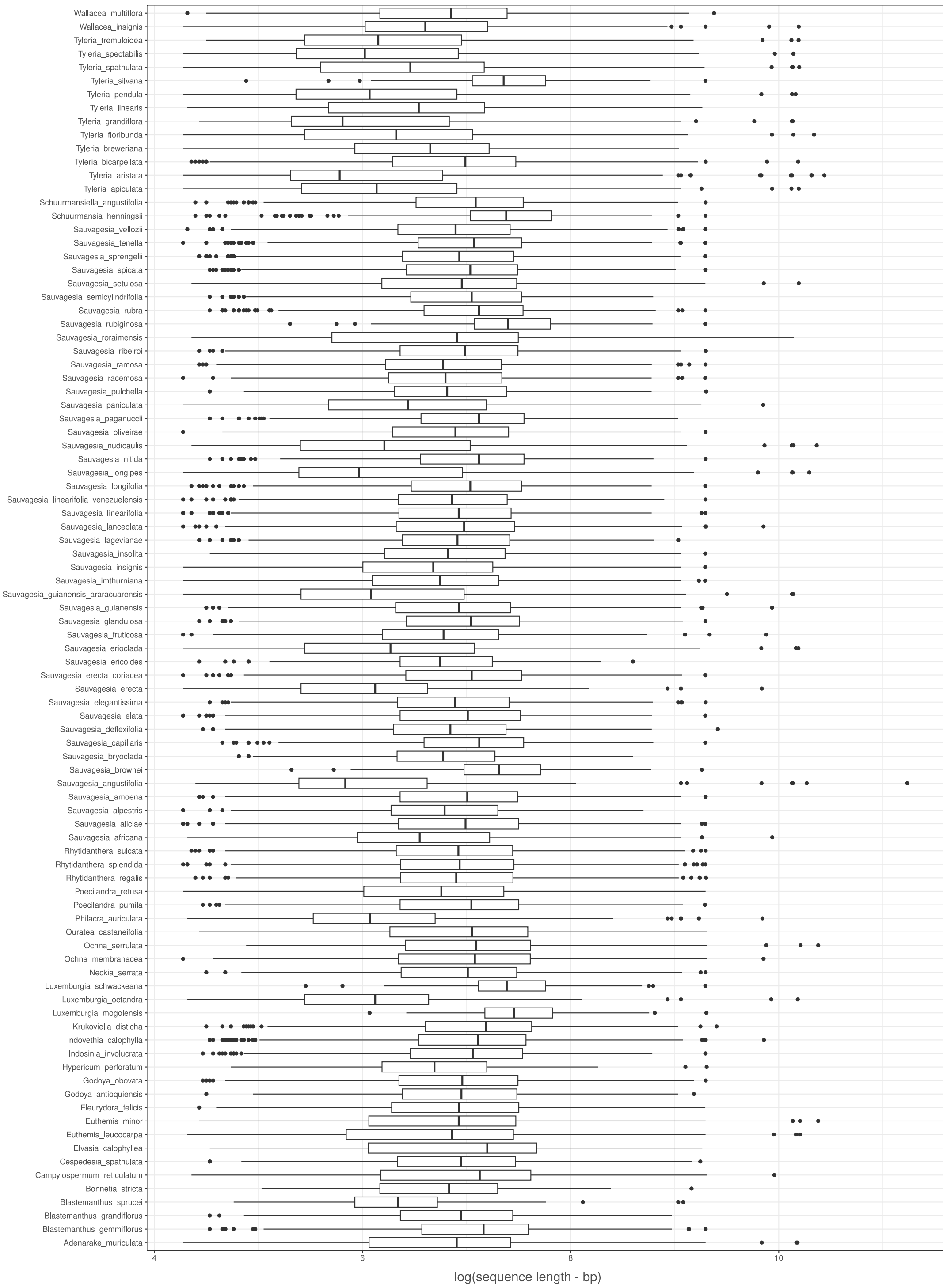


Figure S25: Comparison of the sequence length per gene retrieved by HybPiper for each species.
